# Senescence-associated loss of intestinal α1,2-fucose disrupts a modifiable host-microbiome homeostasis axis in people with HIV

**DOI:** 10.64898/2026.07.09.736798

**Authors:** Leila B. Giron, Maliha W. Shaikh, Thaisa M. Cantu Jungles, Lijuan Zhang, Phillip A. Engen, Nuseybe Bulut, Shalini Singh, Jenna M. Hasson, Ellen Zhang, Shivanjali Shankaran, Corey Neumann, Michelle Villanueva, Alan L. Landay, Thomas J. Hope, Frank J. Palella, Michael J. Corley, Hiroaki Tateno, Bruce Hamaker, Noam Auslander, Ramon Lorenzo Redondo, Ali Keshavarzian, Mohamed Abdel-Mohsen

## Abstract

**Background:** People with HIV (PWH), despite effective antiretroviral therapy (ART), experience disrupted intestinal homeostasis characterized by microbial dysbiosis and impaired intestinal barrier integrity, which contribute to chronic inflammation and aging-associated comorbidities. However, tractable mechanisms contributing to this dysfunction remain poorly defined.

**Objective:** To determine whether acquired loss of intestinal α1,2-fucose, a host-derived intrinsic prebiotic glycan that supports colonization by short-chain fatty acid (SCFA)-producing bacteria essential for intestinal barrier integrity, contributes to microbiome disruption, impaired epithelial resilience, inflammation, and biological aging in PWH.

**Design:** Ileal and colonic biopsies, isolated crypts, stool, and blood from PWH on ART and controls underwent multi-omic analyses. Findings were mechanistically interrogated using stool anaerobic fermentation assays and 3D intestinal organoid models of stress-mediated epithelial disruption.

**Results:** In intestinal tissues, PWH exhibited reduced α1,2-fucosylation and increased senescence-associated expression of the fucose-degrading enzyme α-L-fucosidase. Lower α1,2-fucose tracked with depletion of SCFA-producing bacteria, increased inflammation, and premature biological aging. In anaerobic fermentations, stool from PWH produced fewer SCFAs than controls, whereas supplementation with the human-milk-oligosaccharide-derived α1,2-fucose donor 2′-fucosyllactose restored SCFA production and improved intestinal organoid resilience to stress-mediated disruption.

**Conclusion:** These findings identify acquired loss of intestinal α1,2-fucose as a modifiable host-microbiome mechanism linking epithelial senescence, microbial metabolic dysfunction, impaired barrier resilience, inflammation, and biological aging in treated HIV infection.

**SUMMARY BOX:** *What is already known on this topic:* People with HIV on suppressive antiretroviral therapy frequently have persistent intestinal barrier dysfunction, microbial dysbiosis, chronic inflammation, and accelerated biological aging, but the host mechanisms that maintain this disrupted mucosal state remain incompletely defined.

*What this study adds:* This study identifies acquired loss of intestinal α1,2-fucosylation as a feature of treated HIV infection and links this defect to a host fucosidase-high, senescence-enriched mucosal niche, depletion of SCFA-producing bacteria, impaired tight junction-associated barrier signatures, inflammation, and biological aging phenotypes.

*How this study might affect research, practice or policy:* These findings support intestinal glycan ecology as a modifiable host-microbiome axis and provide a rationale for testing α1,2-fucose-replenishing strategies, such as 2′-fucosyllactose, to restore microbial metabolic output and improve epithelial resilience in people with HIV.

## INTRODUCTION

Despite durable viral suppression on antiretroviral therapy (ART), people with HIV (PWH) exhibit persistent intestinal barrier dysfunction characterized by epithelial injury, increased permeability, and microbial translocation. This disrupted mucosal state contributes to chronic local and systemic inflammation, cellular stress, and aging-associated non-AIDS comorbidities.^1–7^ Hallmarks of this dysfunction include epithelial stress and premature aging programs that reduce mucosal resilience,^8,9^ together with microbial dysbiosis shaped by both HIV-associated factors and sexual practices.^9–19^ This dysbiosis is marked by enrichment of pro-inflammatory taxa and loss of beneficial commensals that normally support intestinal integrity. However, the mechanisms that sustain this chronic microbial dysbiosis and intestinal dysfunction remain poorly defined, limiting efforts to restore intestinal health and reduce inflammation-driven comorbidities in PWH.

One of the most important functional consequences of HIV-associated dysbiosis is the loss of bacteria that produce short-chain fatty acids (SCFAs), including butyrate, acetate, and propionate, which are generated by gut-resident bacteria through carbohydrate fermentation and are essential for intestinal and systemic homeostasis.^20^ Such loss has major biological consequences because SCFAs fuel colonocytes, promote epithelial renewal, strengthen tight junction expression, and restrain inflammatory responses.^21,22^ Thus, depletion of SCFA-producing bacteria may represent a functionally relevant defect that compromises barrier integrity and perpetuates inflammation. Yet the tractable host mechanisms driving the loss of these beneficial bacteria in PWH remain unknown.

An underappreciated candidate mechanism is intestinal glycosylation. Emerging evidence indicates that epithelial glycans help maintain host-microbiota homeostasis by shaping microbial colonization and behavior.^23^ Among these, α1,2-fucose plays a central role.^24^ In the healthy intestine, microbial products activate myeloid cells to produce IL-23, which licenses group 3 innate lymphoid cells (ILC3s) to secrete IL-22. IL-22 engages the IL-22RA1/IL-10RB receptor complex on epithelial cells, thereby inducing the fucosylation machinery through transcriptional and functional activation of FUT2 (α1,2-fucosyltransferase) and other fucosylation enzymes.^25–28^ In addition to epithelial surface glycoproteins, FUT2-dependent α1,2-fucosylation also modifies secreted mucins produced by intestinal goblet cells.^29,30^ Gut bacteria can then cleave and utilize this fucose, generating a host-derived carbohydrate source that preferentially supports beneficial commensals, including *bifidobacteria* and other taxa that sustain SCFA production.^24,26,31–33^ In parallel, fucose can suppress virulence programs in pathobionts through FusKR signaling.^34^ Through these effects, α1,2-fucose helps preserve epithelial barrier integrity and intestinal homeostasis. Conversely, loss of α1,2-fucosylation reduces beneficial bacteria, promotes pathobiont expansion, disrupts the epithelial barrier, and enhances microbial translocation and inflammation.^24,35^

Loss-of-function FUT2 variants in “non-secretors” reduce intestinal α1,2-fucosylation and increase susceptibility to intestinal inflammation, including inflammatory bowel disease.^36–38^ However, whether acquired reductions in intestinal α1,2-fucosylation also occur in chronic conditions associated with intestinal dysfunction, such as chronic HIV infection, and whether such loss contributes to depletion of SCFA-producing bacteria, impaired barrier integrity, inflammation, and premature aging, remains unknown.

Here, we test the hypothesis that chronic HIV infection is associated with loss of intestinal α1,2-fucosylation that acts as a tractable host mechanism contributing to the depletion of SCFA-producing bacteria, microbial dysbiosis, epithelial stress, and disrupted intestinal homeostasis.

Using intestinal tissues, multi-omic profiling, anaerobic fermentations, and intestinal organoid models, we show that PWH exhibit reduced intestinal α1,2-fucosylation associated with senescence-induced expression of the fucose-degrading enzyme α-L-fucosidase. Consistent with the established role of α1,2-fucose in maintaining intestinal homeostasis by supporting colonization of SCFA-producing bacteria, depletion of α1,2-fucose in PWH was associated with loss of SCFA-producing bacteria, inflammation, and biological aging phenotypes. We further show that supplementation with the human-milk-derived α1,2-fucose donor 2′-fucosyllactose (2’FL) restores SCFA production and improves organoid resilience to stress-mediated disruption, identifying intestinal α1,2-fucosylation as a modifiable host-microbiome axis in ART-treated PWH.

## RESULTS

### PWH on ART exhibit reduced intestinal α1,2-fucosylation despite similar FUT2 secretor genotype

To examine the impact of chronic treated HIV infection on intestinal glycosylation, we collected ileal and colonic biopsies, as well as stool and blood, from 25 PWH on ART and 22 matched people without HIV (PWoH) (**Table S1**). Using these samples, we performed a multi-omic investigation to determine whether PWH exhibit depletion of intestinal α1,2-fucose which could contribute to the persistent intestinal dysfunction observed despite long-term ART (**Figure 1A**).

**Figure 1.**
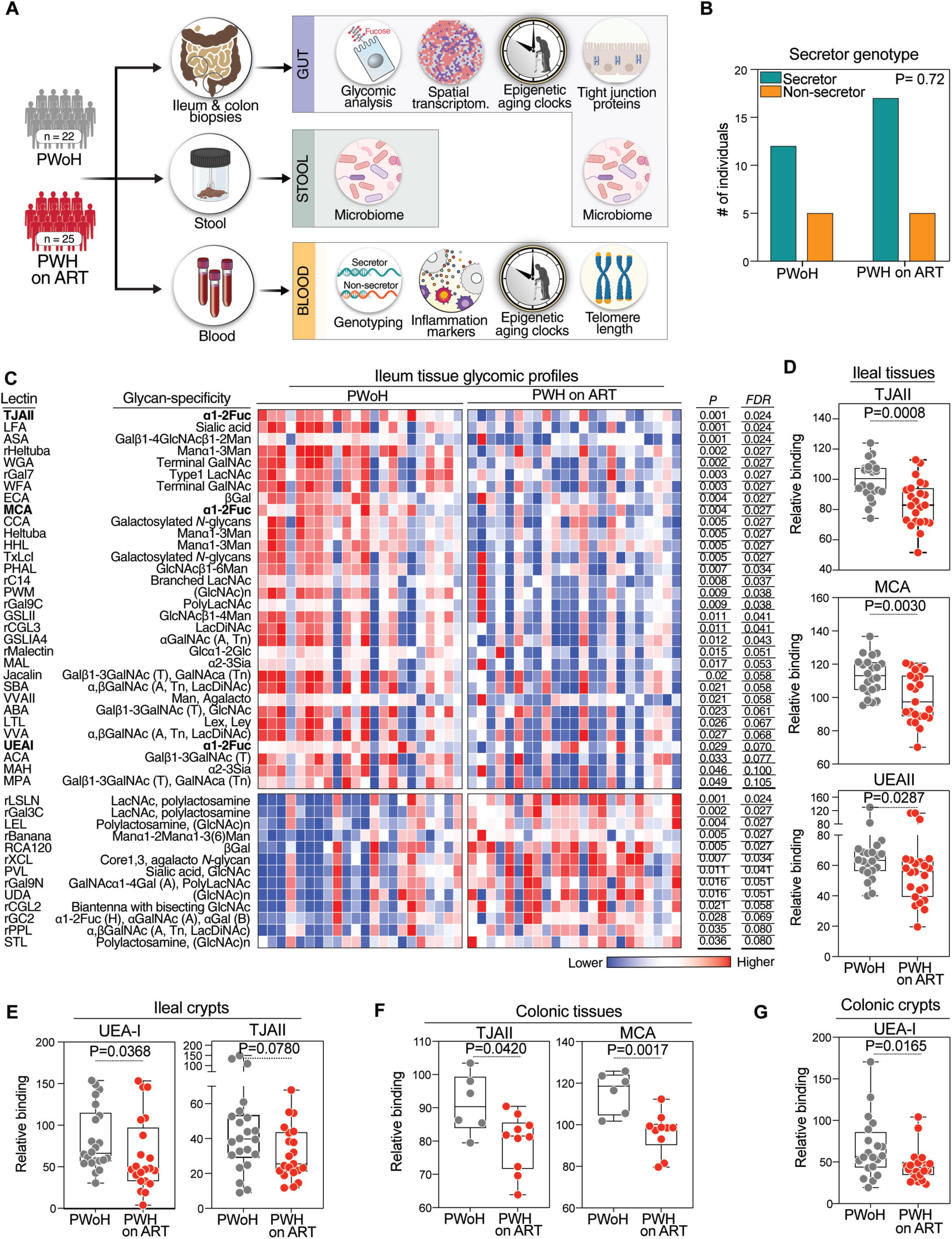
PWH on ART exhibit reduced intestinal α 1,2-fucosylation despite similar FUT2 secretor genotype. **(A)** Overview of the study design. Ileal and colonic biopsies, isolated intestinal crypts, stool, and blood were collected from PWH on ART and matched PWoH for integrated glycomic, genomic, microbiome, inflammatory, biological aging, fermentation, and organoid analyses. **(B)** Frequency of FUT2 secretor and non-secretor genotypes in PWH on ART and PWoH. Data were analyzed using a two-sided Fisher’s exact test. DNA samples from n= 22 PWH and n= 17 PWoH. **(C)** Lectin microarray analysis of whole ileal and colonic tissues showing global intestinal glycomic differences between PWH on ART and PWoH. Lectins recognizing α1,2-fucose, including TJA-II, UEA-I, and MCA, are highlighted. Red indicates higher expression, whereas blue indicates lower expression. Data were analyzed using two-sided Mann-Whitney U tests. **(D-G)** Quantification of α1,2-fucose-recognizing lectin signals in whole ileal tissue n=23 PWH and n=22 PWoH), whole colonic tissue (n=10 PWH and n=6 PWoH), isolated ileal crypts (n=21 PWH and n=22 PWoH), and isolated colonic crypts (n=23 PWH and n=20 PWoH) from PWH on ART and PWoH. Box-and-whisker plots show all data points. Data were analyzed using two-sided Mann-Whitney U tests.

Because loss of α1,2-fucosylation can result from FUT2 non-secretor status, we first assessed FUT2 secretor genotype by genotyping the rs516246 allele and found no difference in the frequency of the non-secretor genotype between PWH on ART and PWoH (**Figure 1B**). We next performed glycomic analysis of total ileal and colonic tissues, as well as isolated crypts from both sites, using lectin microarray technology.^39–46^ This platform employs immobilized lectins with defined glycan-binding specificities (**Table S2**), and the degree of binding to each lectin reflects the abundance of the corresponding lectin-binding glycans in a given sample. As shown in **Figure 1C** and **Figure S1**, ileal and colonic biopsies from PWH on ART exhibited multiple glycomic alterations relative to PWoH, including reduced binding to lectins specific for α1,2-fucose, including TJA-II, UEA-I, and MCA. Importantly, this reduction was consistently observed in both whole tissues and isolated crypts from the ileum and colon (**Figure 1D-G**), indicating that the defect was not restricted to one intestinal site and was evident within the epithelial crypt niche itself. Together, these data show that PWH on ART exhibit a reduction in intestinal α1,2-fucosylation that is not explained by differences in FUT2 secretor genotype, supporting an acquired defect in mucosal fucosylation during treated HIV infection.

### Reduced intestinal α1,2-fucosylation in PWH is linked to a host fucosidase-high senescent mucosal niche rather than increased microbial fucosidase capacity

After identifying reduced intestinal α1,2-fucosylation in PWH on ART, we next asked whether this defect could reflect increased fucose degradation. Because fucose can be released from host glycans by microbial fucosidases, we first tested whether the gut microbiome of PWH exhibits increased bacterial α-fucosidase capacity. To address this, we analyzed stool shotgun metagenomic data for bacterial α-fucosidase-associated genes and taxa. This analysis did not support a model in which increased microbial fucosidase activity explains the intestinal α1,2-fucose defect in PWH. No bacterial α-fucosidase-associated feature showed a significant increase in PWH after correction for multiple testing. In fact, when nominal comparisons were examined, microbial α-fucosidase-associated features tended to be lower, not higher, in PWH on ART relative to PWoH (**Figure 2A**). These findings argue against increased bacterial fucose degradation as a mechanism underlying intestinal α1,2-fucose depletion in treated HIV infection. Instead, the reduction in microbial fucosidase-associated features may be consistent with reduced microbial access to, or demand for, host-derived fucose once mucosal α1,2-fucose availability is diminished.

**Figure 2.**
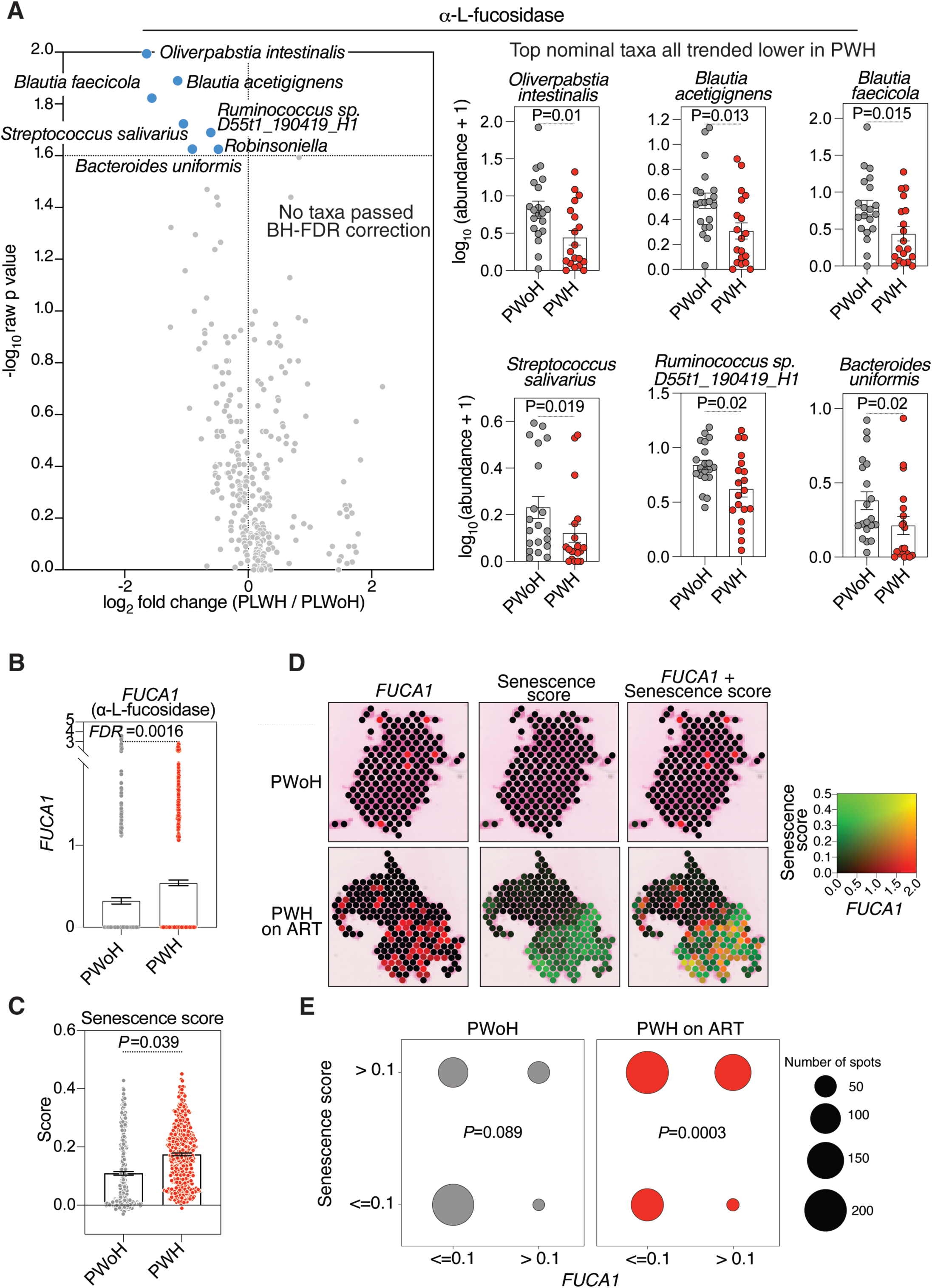
Reduced intestinal α1,2-fucosylation in PWH is linked to a host FUCA1-high senescent mucosal niche rather than increased microbial fucosidase capacity. **(A)** Shotgun metagenomic analysis of stool samples assessing bacterial α-fucosidase-associated genes and taxa in PWH on ART and PWoH. No microbial α-fucosidase-associated features were significantly increased in PWH after correction for multiple testing; nominal two-sided Mann-Whitney U tests trended toward reduced microbial fucosidase-associated capacity in PWH. Fecal samples from 19 PWH and 20 PWoH. **(B)** Spatial transcriptomic analysis showing increased expression of the host fucose-degrading enzyme FUCA1 in intestinal tissues from PWH on ART compared with PWoH. Differential expression of FUCA1 between groups was assessed with donor identity included as a latent variable; the reported FDR represents the Benjamini-Hochberg-adjusted p value from this analysis. **(C)** Quantification of senescence-high spatial regions in intestinal tissues from PWH on ART and PWoH based on a previously described senescence score. Indicated p value reflects the contrast in senescence scores between Visium slide spots with tissue from PWH on ART and PWoH, derived from the robust linear model. **(D-E)** Spatial mapping of FUCA1-high regions and senescence-enriched regions in representative intestinal tissues from PWH on ART and PWoH, showing increased overlap between FUCA1 expression and senescence-enriched mucosal niches in PWH. Ileal tissues from two PWH and two PWoH. Reported p-values reflect pairwise contrasts between FUCA1-positive and FUCA1-negative spots within each HIV status group, derived from the estimated marginal means of the binomial GLM and adjusted for multiple comparisons.

Because the microbial fucosidase analysis did not explain the observed mucosal fucose defect, we next investigated whether host fucose degradation contributes to intestinal α1,2-fucose loss in PWH. We focused on FUCA1 (α-L-fucosidase 1), which encodes lysosomal α-L-fucosidase, a host glycosidase that removes terminal fucose residues from glycoconjugates. FUCA1 was of particular interest because senescent cells undergo lysosomal remodeling and increased lysosomal hydrolase activity, and α-L-fucosidase has been identified as a senescence-associated lysosomal enzyme.^47–51^ Using spatial transcriptomic profiling of intestinal tissues, we found that intestinal tissues from PWH on ART exhibited significantly higher FUCA1 expression than PWoH (**Figure 2B**). To determine whether FUCA1 expression was linked to senescence programs in intestinal tissues, we calculated a senescence score, as previously described,^52^ and identified regions with high senescence enrichment. Intestinal tissues from PWH contained significantly more senescence-high regions than tissues from PWoH (**Figure 2C**). Spatial mapping further showed that FUCA1-high regions in PWH overlapped with senescence-enriched niches, whereas this pattern was much less evident in tissues from PWoH (**Figure 2D-E**). Together, these data indicate that the intestinal α1,2-fucose defect in PWH is not explained by increased microbial fucosidase capacity but is instead linked to a host FUCA1-high, senescence-enriched mucosal niche.

### Senescence shifts intestinal epithelial fucose homeostasis toward degradation despite preserved production signaling

To further define the host program linked to increased intestinal fucose degradation, we next examined transcriptional pathways associated with the FUCA1-high mucosal niche. Intestinal tissues from PWH exhibited increased expression of senescence- and senescence-associated secretory phenotype (SASP)-related genes, including MMP9, IGFBP3, ATM, MAP3, IGFBP7, and CDKN1A (**Figure 3A**). These genes are consistent with established senescence programs involving DNA damage signaling, CDKN1A/p21-mediated cell-cycle arrest, matrix remodeling, and SASP-associated secreted factors that can contribute to tissue dysfunction and chronic inflammation.

**Figure 3.**
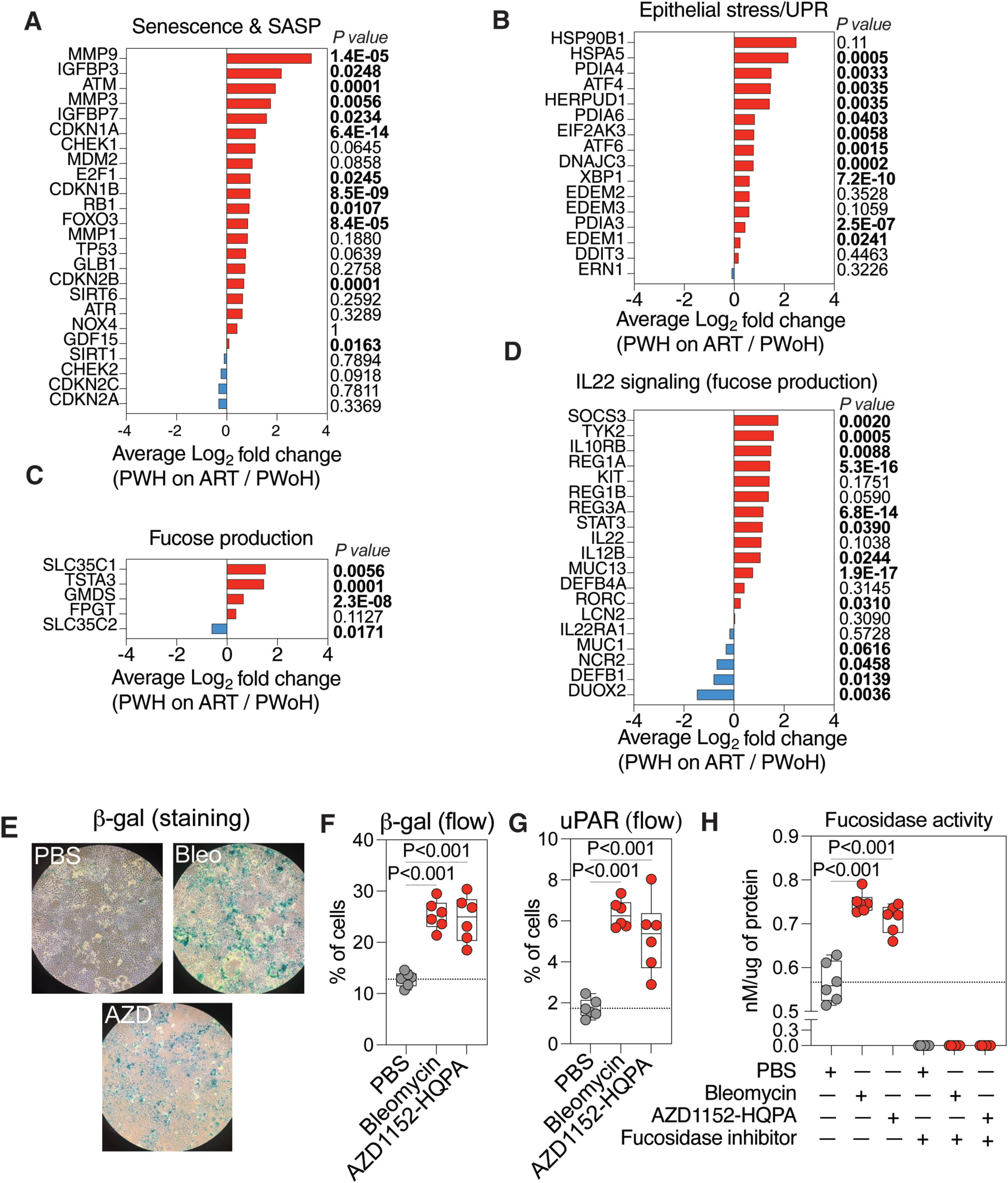
Senescence shifts intestinal epithelial fucose homeostasis toward degradation despite preserved production signaling. **(A)** Expression of senescence- and senescence-associated secretory phenotype-related genes in intestinal tissues from PWH on ART and PWoH. **(B)** Expression of epithelial stress- and unfolded protein response-related genes. **(C)** Expression of genes involved in GDP-fucose biosynthesis and fucose transport. **(D)** Expression of genes involved in the IL-22 signaling axis. P values in panels A-D were derived from the spatial transcriptomics Differential Expression Analysis performed between groups with donor identity included as a latent variable. Ileal tissues from two PWH and two PWoH. **(E)** Representative senescence-associated β-galactosidase staining of Caco-2 cells treated with vehicle control or senescence-inducing agents. **(F)** Flow cytometric quantification of β-galactosidase-positive Caco-2 cells following chemical induction of senescence. **(G)** uPAR expression in control and senescent Caco-2 cells. **(H)** Active α-L-fucosidase activity measured using a fucose-cleavage assay in control and senescent Caco-2 cells, with a fucosidase inhibitor control confirming assay specificity. Data in panels F-H are shown as box-and-whisker plots with all data points displayed (n=6 biological replicates). Analyses in panels F-H were performed using one-way ANOVA, with multiple comparisons corrected using the two-stage step-up method of Benjamini, Krieger, and Yekutieli.

In parallel, epithelial stress and unfolded protein response programs were enriched in PWH tissues, as reflected by increased expression of HSPA5, PDIA4, ATF4, HERPUD1, EIF2AK3, and XBP1 (**Figure 3B**). These genes map onto canonical epithelial stress and endoplasmic reticulum stress pathways, including the PERK-eIF2α-ATF4 and IRE1-XBP1 arms of the unfolded protein response. Thus, the FUCA1-high intestinal niche in PWH is embedded within a broader tissue state characterized by senescence, epithelial stress, and mucosal dysfunction.

We next asked whether the observed loss of α1,2-fucose might instead reflect impaired fucose production. Unexpectedly, genes linked to epithelial fucose biosynthesis and transport were not globally reduced in PWH. Rather, transcripts involved in GDP-fucose synthesis and transport, including TSTA3, GMDS, SLC35C1, and related fucose-production genes, were preserved or increased in PWH tissues (**Figure 3C**). Similarly, genes in the IL-22 signaling axis, a recognized upstream regulator of epithelial fucosylation, were also preserved or increased (**Figure 3D**).^25–28^ These findings argue against a simple failure of the epithelial fucosylation machinery and instead support a model in which intestinal α1,2-fucose depletion in PWH is linked predominantly to enhanced host fucose degradation within senescence- and stress-enriched mucosal niches.

Although the tissue analyses linked FUCA1 expression to senescence-associated intestinal regions *in vivo*, they did not establish whether senescence is sufficient to directly increase epithelial fucosidase activity. We therefore tested this question in an intestinal epithelial cell model. Chemical induction of senescence in Caco-2 cells increased markers of senescence, including β-galactosidase (β-gal), as evident by staining (**Figure 3E**) and flow cytometric analysis (**Figure 3F**), relative to non-senescent controls. In addition, these senescent cells exhibited increased expression of other senescence markers, including uPAR^53,54^ (**Figure 3G**). These senescent Caco-2 cells exhibited higher active α-L-fucosidase activity, measured by a fucose-cleavage assay, than control cells (**Figure 3H**). These findings provide direct functional evidence that senescence is sufficient to induce epithelial fucosidase activity, extending prior work identifying α-L-fucosidase as a senescence-associated lysosomal feature.^47–51^ Taken together, data in Figures 2 and 3 support a model in which the intestinal α1,2-fucose defect in PWH on ART is not driven by increased microbial fucosidase capacity, but instead by a host senescence-associated fucose-degradation program marked by increased FUCA1 expression and α-L-fucosidase activity.

### Loss of intestinal α1,2-fucose is associated with inflammation and biological aging phenotypes

Because epithelial α1,2-fucosylation is an important component of intestinal homeostasis, we next asked whether depletion of intestinal α1,2-fucose in PWH is linked to inflammatory and aging-related phenotypes. Across intestinal compartments, lower α1,2-fucose levels correlated with multiple plasma markers linked to microbial translocation, epithelial injury, and myeloid inflammation, including LPS-binding protein (LBP), intestinal fatty-acid-binding protein (IFABP), sCD14, sCD163, CXCL9, IL-12p70, MIP-1α, and MPO (**Figure 4A**). These associations were especially prominent in colonic tissues and crypts, where reduced binding to α1,2-fucose-recognizing lectins tracked with higher inflammatory burden.

**Figure 4.**
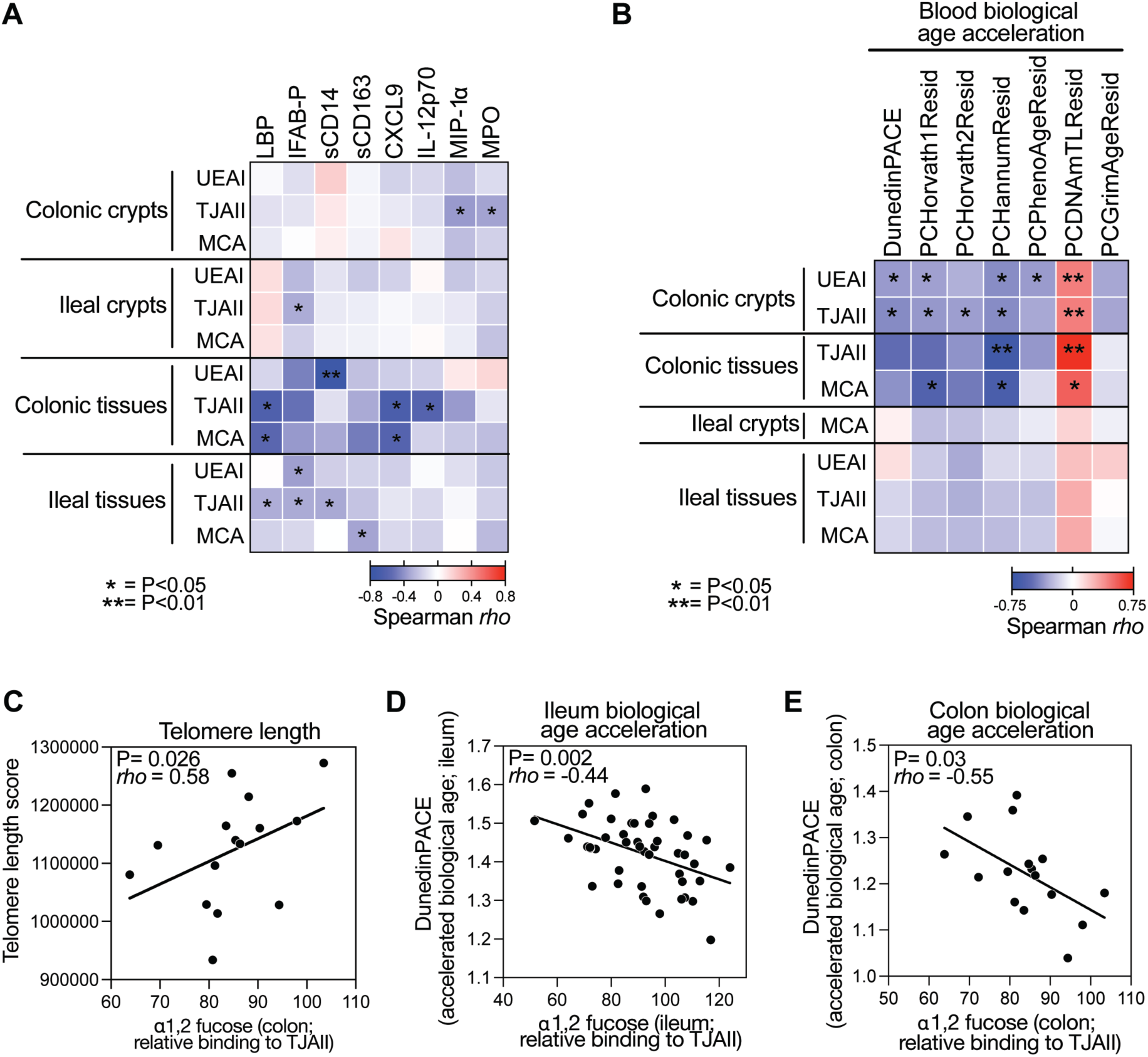
Loss of intestinal α1,2-fucose is associated with inflammation and biological aging phenotypes. **(A)** Spearman’s rank correlation heatmap showing associations between intestinal α1,2-fucose-recognizing lectin signals and plasma markers of microbial translocation, epithelial injury, myeloid activation, and inflammation, including LBP, IFABP, sCD14, sCD163, CXCL9, IL-12p70, MIP-1α, and MPO. Red indicates positive correlation, whereas blue indicates negative correlation. **(B)** Spearman’s rank correlation heatmap showing associations between intestinal α1,2-fucose levels and systemic biological age acceleration measured in blood using multiple epigenetic aging clocks. Red indicates positive correlation, whereas blue indicates negative correlation. **(C)** Spearman’s rank association between colonic α1,2-fucose levels and telomere length score measured in PBMCs (n=15 samples). **(D-E)** Spearman’s rank associations between ileal (n=45 samples) and colonic (n=16 samples) α1,2-fucose levels and intestinal biological age acceleration measured by DunedinPACE.

We next examined whether intestinal α1,2-fucose levels were linked to biological aging measures. Lower α1,2-fucose levels were broadly associated with greater systemic biological age acceleration measured in blood across multiple epigenetic aging clocks, with the strongest signals observed in colonic crypts and colonic tissues (**Figure 4B**). Higher colonic α1,2-fucose was also positively associated with telomere length score measured in PBMCs (**Figure 4C**), whereas lower ileal and colonic α1,2-fucose levels were associated with greater intestinal biological age acceleration measured by DunedinPACE (**Figure 4D-E**). Together, these findings link the intestinal α1,2-fucose defect in treated HIV infection to a more inflamed mucosal and systemic milieu and to both intestinal and systemic biological aging phenotypes.

### Reduced intestinal α1,2-fucose is linked to depletion of SCFA-producing bacteria, lower microbiome diversity, and impaired barrier-supportive signatures

To understand how loss of intestinal α1,2-fucose might contribute to inflammation and biological aging, we next examined its relationship to the gut microbiome. This was biologically plausible because host-derived fucose can function as a nutrient source and ecological cue for commensal bacteria, helping to shape microbial colonization, microbial metabolism, and host-microbiota symbiosis.^24,26,31–33^ We first asked whether reduced intestinal α1,2-fucose is linked to loss of SCFA-producing bacteria. Both ileal and colonic microbiomes from PWH on ART exhibited depletion of multiple SCFA-producing taxa relative to PWoH (**Figure 5A**). These taxa included members of *Alistipes*, *Butyricimonas*, *Odoribacter*, *Coprococcus*, *Subdoligranulum*, *Fusicatenibacter*, *Faecalibacterium*, *Agathobacter*, and *Butyricicoccus*, among others.

**Figure 5.**
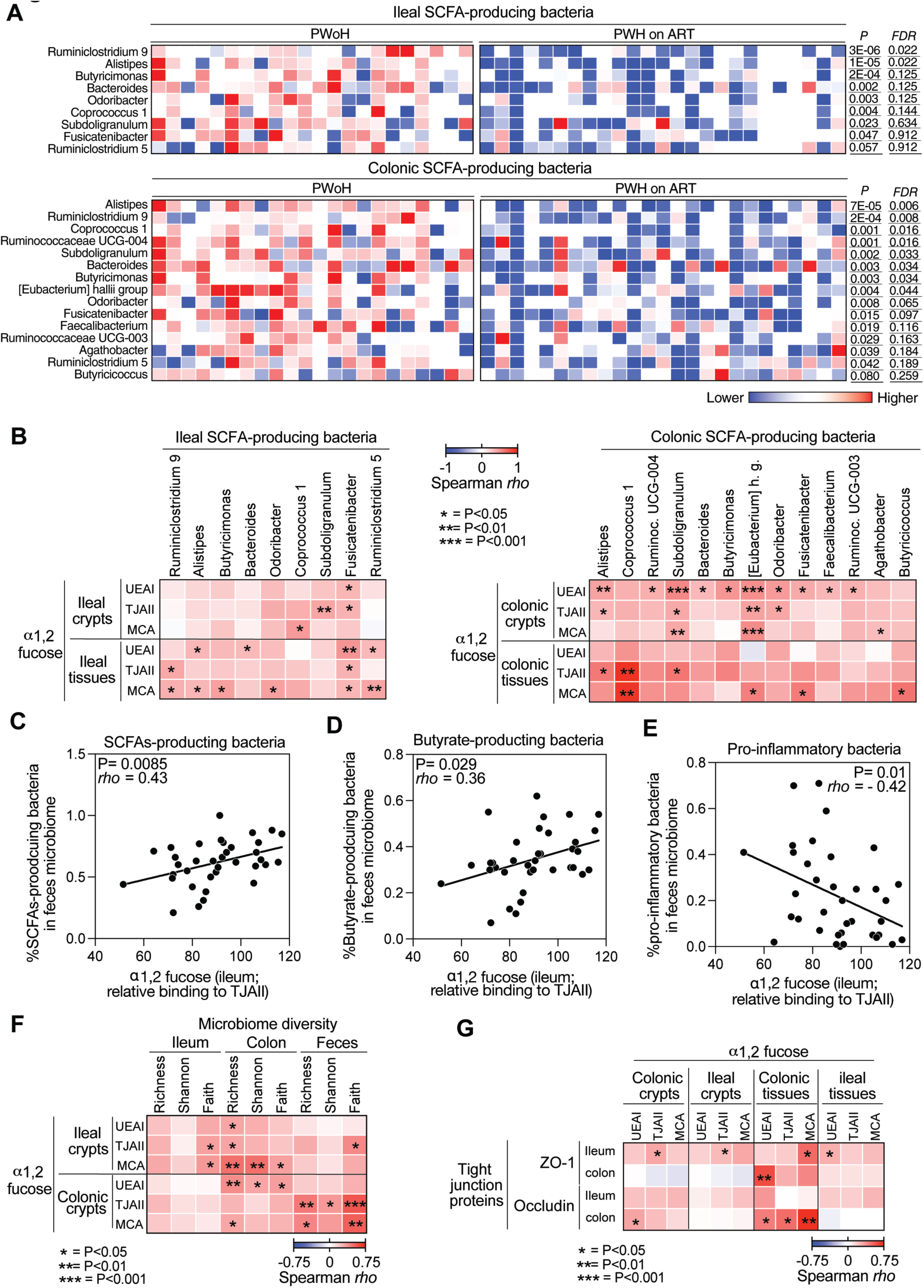
Reduced intestinal α1,2-fucose is linked to depletion of SCFA-producing bacteria, lower microbiome diversity, and impaired barrier-supportive signatures. **(A)** Differential abundance of short-chain fatty acid (SCFA)-producing bacterial taxa in ileal and colonic microbiomes from PWH on ART and PWoH. Taxa shown include members of *Alistipes*, *Butyricimonas*, *Odoribacter*, *Coprococcus*, *Subdoligranulum*, *Fusicatenibacter*, *Faecalibacterium*, *Agathobacter*, *<u>Butyricicoccus</u>*, and related SCFA-producing groups. Two-sided Mann-Whitney U tests corrected using were calculated using the Benjamini-Hochberg procedure. Samples from n=25 PWH and n=22 PWoH. **(B)** Spearman’s rank correlation analysis between intestinal α1,2-fucose levels and SCFA-producing bacterial taxa across ileal and colonic compartments. Red indicates positive correlation, whereas blue indicates negative correlation. **(G)** Spearman’s rank association between ileal α1,2-fucose levels and the overall proportion of fecal SCFA-producing bacteria. **(D)** Spearman’s rank association between ileal α1,2-fucose levels and the proportion of fecal butyrate-producing bacteria. **(E)** Spearman’s rank association between ileal α1,2-fucose levels and the abundance of putatively pro-inflammatory bacteria. N= 36 samples. **(F)** Spearman’s rank correlation analysis between intestinal α1,2-fucose levels and microbiome diversity metrics, including richness, Shannon diversity, and Faith phylogenetic diversity. Red indicates positive correlation, whereas blue indicates negative correlation. **(G)** Spearman’s rank correlation analysis between intestinal α1,2-fucose levels and epithelial tight junction proteins ZO-1 and occludin measured by immunohistochemistry. Red indicates positive correlation, whereas blue indicates negative correlation.

Consistent with a functional link between mucosal α1,2-fucose and microbial metabolism, intestinal α1,2-fucose levels positively correlated with the abundance of multiple SCFA-producing taxa in both ileal and colonic compartments (**Figure 5B**). At the community level, we calculated the relative abundance of SCFA-producing bacteria, butyrate-producing bacteria, and pro-inflammatory bacteria (**Table S3**). Higher ileal α1,2-fucose correlated with a greater overall proportion of fecal SCFA-producing bacteria (**Figure 5C**) and butyrate-producing bacteria (**Figure 5D**), whereas lower ileal α1,2-fucose correlated with increased abundance of putatively pro-inflammatory bacteria (**Figure 5E**). These findings suggest that depletion of intestinal α1,2-fucose is linked not only to loss of beneficial fermenters but also to a broader shift toward a more pro-inflammatory microbiome configuration. This relationship extended beyond individual taxa. Across intestinal and fecal compartments, higher α1,2-fucose levels were positively associated with multiple measures of microbiome diversity, including richness, Shannon diversity, and Faith phylogenetic diversity (**Figure 5F**). We next examined whether α1,2-fucose levels were linked to intestinal barrier integrity, estimated by immunohistochemical measurement of tight junction proteins (**Figure S2**). Higher α1,2-fucose levels correlated with increased expression of the epithelial tight junction proteins ZO-1 and occludin, whereas depletion of α1,2-fucose correlated with lower levels of these proteins, particularly in the colon (**Figure 5G**). These findings are consistent with the known role of SCFA-producing communities, especially butyrate-producing communities, in supporting epithelial energetics, tight junction maintenance, and barrier function. Together, these results support a model in which loss of intestinal α1,2-fucose is linked to depletion of SCFA-producing bacteria, reduced microbial community diversity, and impaired epithelial barrier-supportive programs.

### 2’FL restores microbial SCFA production in stool from PWH on ART

We next asked whether the α1,2-fucose defect in PWH has functional consequences for microbial metabolic output and whether these effects can be rescued by fucose replenishment. Human milk oligosaccharides (HMOs), complex carbohydrates that are indigestible by the infant intestine,^55^ are a naturally abundant source of α1,2-fucose. Rather than directly nourishing the infant, HMOs shape and support the infant gut microbiota, promoting the growth of beneficial SCFA-producing bacteria while suppressing pathobionts.^56^ This functional specialization is a key reason why breastfed infants harbor distinct, and often more beneficial, microbial profiles than formula-fed infants.^57^ Among HMOs, 2′FL, which consists of α1,2-fucose linked to lactose, is highly abundant.^58^ Given its established safety profile, as 2′FL is already incorporated into infant formulas and is safe in adults,^59,60^ we hypothesized that this naturally occurring glycan with an established safety profile could be repurposed to replenish depleted α1,2-fucose in PWH and restore healthy microbiota function.

We therefore performed anaerobic stool fermentation assays using stool from PWoH and PWH on ART, with or without supplementation with 2′FL (**Figure 6A**). These two groups had similar chronological age (**Figure 6B**), but PWH on ART exhibited accelerated biological aging, as estimated by the residual of the Horvath epigenetic clock of biological aging^61^ (**Figure 6C**), and higher levels of inflammatory aging^62^, as estimated by CXCL9 levels (**Figure 6D**). Relative to stool from PWoH, stool from PWH on ART produced significantly lower levels of SCFAs (**Figure 6E**). Importantly, supplementation of stool from PWH with 2′FL significantly increased SCFA production, including propionate, butyrate, and acetate (**Figure 6F**). The capacity of each stool sample to produce SCFAs in response to fermentation was inversely correlated with biological age measured by DNA methylation clocks and with inflammatory markers such as CXCL9 (**Figure 6G**). These findings indicate that although stool from PWH on ART exhibits reduced basal SCFA output, the microbiota retains the capacity to respond metabolically to α1,2-fucose replenishment.

**Figure 6.**
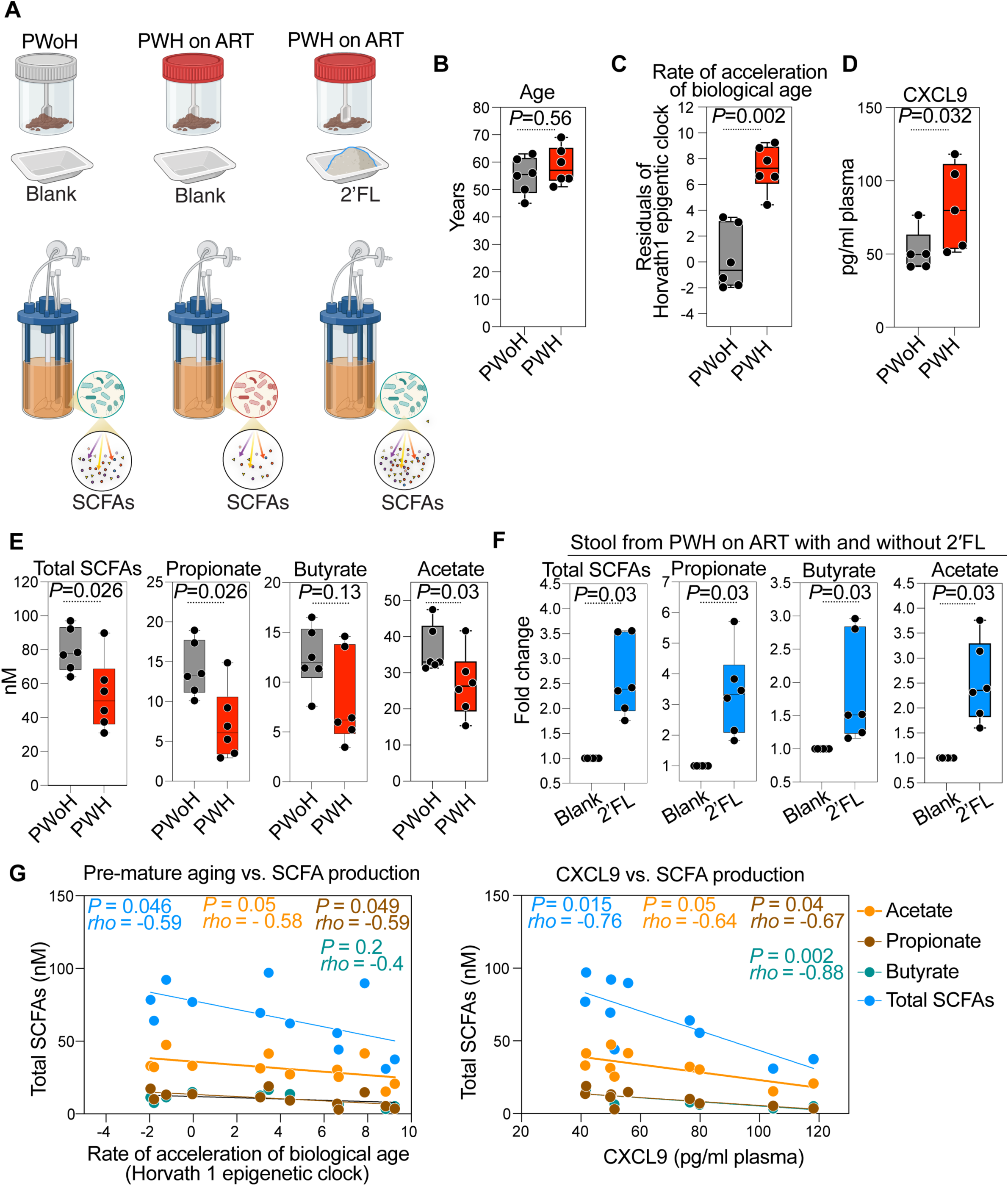
2′FL restores microbial SCFA production in stool from PWH on ART. **(A)**Schematic of the anaerobic stool fermentation assay. Stool samples from six PWH on ART and six PWoH were fermented under anaerobic conditions with or without supplementation with 2′FL, and SCFA production was quantified. **(B)** Chronological age of PWH on ART and PWoH included in the fermentation experiments. **(C)** Biological age acceleration estimated using residuals from the Horvath epigenetic clock. **(D)** CXCL9 levels as a marker of inflammatory aging. **(E)** SCFA production by stool from PWH on ART and PWoH under basal fermentation conditions. Data in panels B-E are shown as box-and-whisker plots show all data points. Data were analyzed using two-sided Mann-Whitney U tests. **(F)** SCFA production by stool from PWH on ART following supplementation with 2′FL, including total SCFAs, propionate, butyrate, and acetate. Wilcoxon tests. **(G)** Spearman’s rank associations between fermentation-induced SCFA production capacity and biological aging or inflammatory markers, including DNA methylation age measures and CXCL9.

### 2′FL-conditioned microbial metabolites improve epithelial resilience to inflammatory and ethanol-mediated disruption

We next tested whether the metabolic effects of 2′FL supplementation translate into improved epithelial resilience. To do this, we exposed colon organoids to fermentation supernatants generated from stool of PWoH or PWH on ART, with or without 2′FL supplementation, and then challenged the organoids with either TNFα plus IFNγ or ethanol. Organoid permeability was assessed using a FITC-dextran intestinal permeability assay.^9^ Increased FITC-dextran fluorescence intensity was used as a readout of reduced organoid integrity and increased permeability. Under inflammatory or ethanol-mediated stress, supernatants derived from stool of PWH without 2′FL were associated with greater epithelial disruption. In contrast, supernatants generated in the presence of 2′FL significantly improved organoid resilience to both cytokine-mediated and ethanol-mediated injury (**Figure 7A-B**). These findings indicate that 2′FL does not merely increase microbial SCFA production in isolation but also generates fermentation products that improve epithelial resistance to stress-mediated barrier injury. This functional rescue is consistent with the established role of SCFAs in supporting epithelial barrier integrity and restraining inflammatory damage.^9–19^

**Figure 7.**
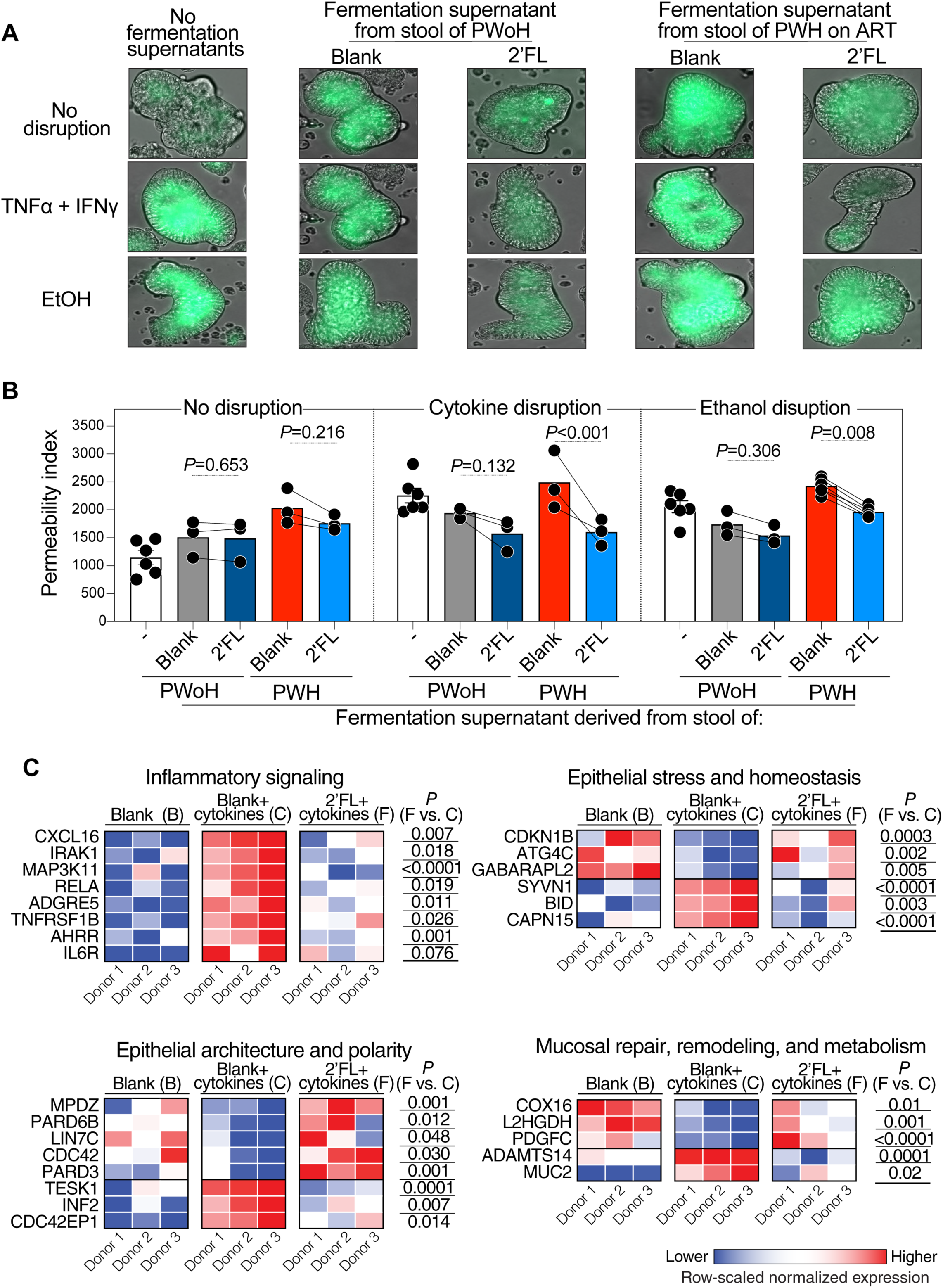
2′FL-conditioned microbial metabolites improve epithelial resilience and restore epithelial homeostasis programs in intestinal organoids. **(A)** Experimental design for testing the epithelial effects of fermentation supernatants. Colon organoids were exposed to supernatants generated from anaerobic stool fermentations using stool from n=3-6 PWH on ART or n=3 PWoH, with or without 2′FL supplementation, and then challenged with TNFα plus IFNγ or ethanol. Organoid permeability was measured using FITC-dextran fluorescence. **(B)** Quantification of FITC-dextran permeability in colon organoids exposed to fermentation supernatants under cytokine- or ethanol-mediated stress. Increased FITC-dextran fluorescence indicates reduced organoid integrity and increased permeability. Data were analyzed using mixed effects analyses corrected with using the two-stage step-up method of Benjamini, Krieger, and Yekutieli. **(C)** RNA-seq analysis of cytokine-exposed organoids treated with fermentation supernatants from PWH stool generated with or without 2′FL. Red indicates higher expression, whereas blue indicates lower expression. 2′FL-conditioned supernatants reduced inflammatory and stress-associated epithelial signatures and shifted epithelial organization and repair-related genes toward a more resilient state, consistent with the functional organoid rescue. P values were calculated using a paired Wald test in DESeq2. The full statistical analysis for this panel is shown in Figure S3.

To define the epithelial transcriptional programs associated with improved organoid resilience, we performed RNA-seq on organoids exposed to fermentation supernatants generated from stool of PWH on ART with or without 2′FL supplementation in the presence of cytokine disruption. Transcriptomic analysis showed that, in cytokine-exposed organoids, 2′FL shifted a selected set of gut-relevant genes toward a more resilient epithelial state, characterized by attenuation of cytokine and inflammatory signaling genes, including *CXCL16, IRAK1, MAP3K11, RELA, TNFRSF1B, AHRR, and IL6R*; partial normalization of epithelial stress and homeostasis genes, including *CDKN1B, ATG4C, GABARAPL2, SYVN1, BID, and CAPN15*; and favorable shifts in genes linked to epithelial architecture, polarity, and cytoskeletal organization, including *PARD3, PARD6B, MPDZ, CDC42, LIN7C, TESK1, INF2, and CDC42EP1*. Additional changes in *COX16, L2HGDH, PDGFC, ADAMTS14, and MUC2* suggest effects on mucosal repair, remodeling, and metabolism (**Figure 7C**, full statistics in **Figure S3**). Together, these data support a model in which 2′FL does not simply increase canonical barrier gene expression but instead promotes a broader epithelial resilience program marked by reduced inflammatory stress, improved epithelial organization, and partial restoration of homeostatic and repair-associated pathways. Together with the fermentation and organoid data, these findings support a model in which α1,2-fucose replenishment restores microbial metabolic outputs that secondarily promote epithelial homeostasis and resistance to stress-mediated disruption.

Collectively, our data support a model (**Figure 8**) in which chronic treated HIV infection is associated with an acquired loss of intestinal α1,2-fucosylation despite similar FUT2 secretor genotype. This defect is not explained by increased microbial fucosidase capacity. Instead, it is linked to a host senescence-associated fucose-degradation program marked by increased FUCA1 expression and α-L-fucosidase activity. Loss of intestinal α1,2-fucose is associated with depletion of SCFA-producing bacteria, reduced microbiome diversity, impaired tight junction expression, increased inflammation, and biological aging phenotypes. Replenishment of α1,2-fucose with 2′FL restores microbial SCFA production, improves epithelial resilience, and promotes epithelial homeostasis programs. These findings identify intestinal α1,2-fucosylation as a modifiable host-microbiome axis that links epithelial stress, microbial metabolism, inflammation, and barrier dysfunction in treated HIV infection.

**Figure 8.**
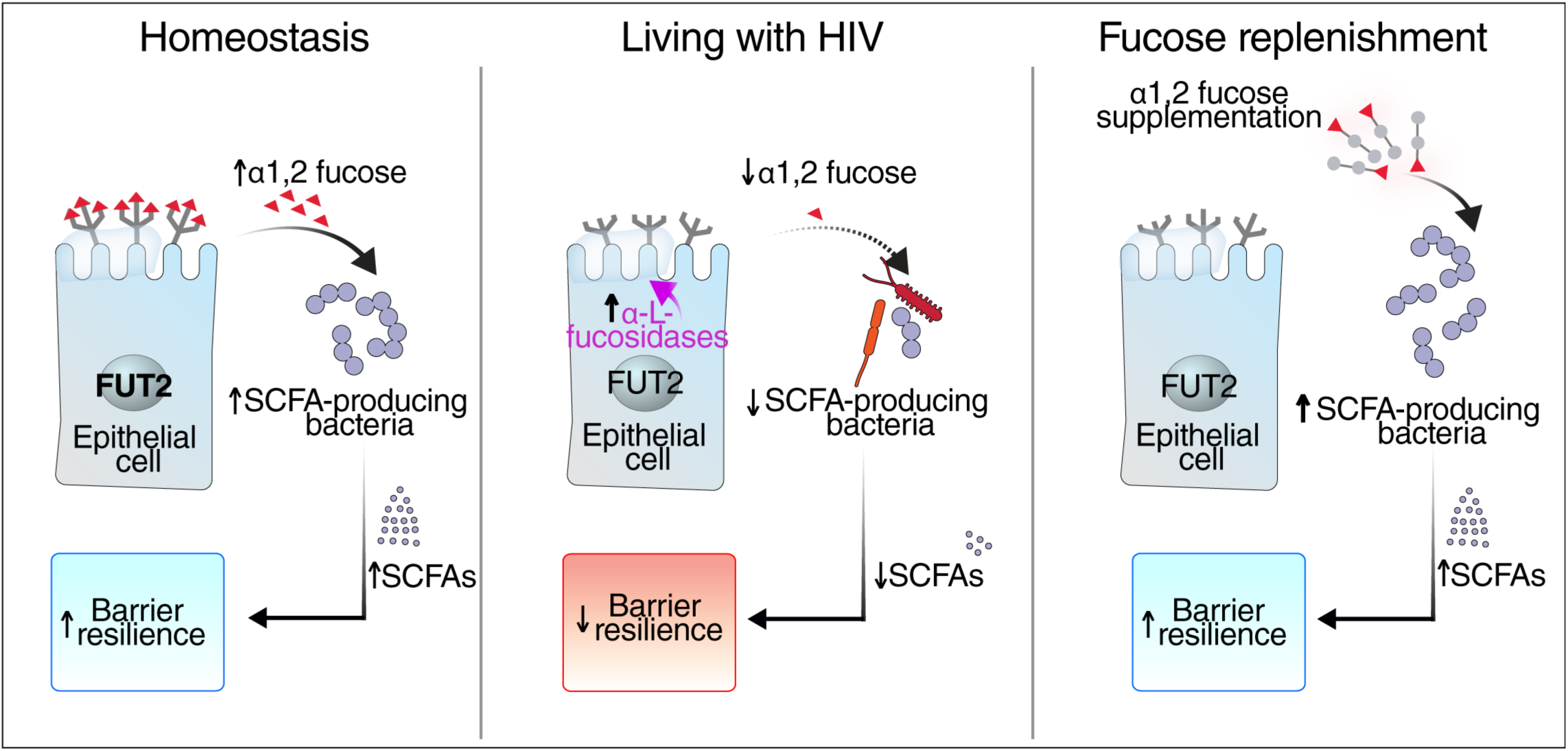
A model of intestinal α1,2-fucose loss and restoration in treated HIV infection. In intestinal homeostasis, α1,2-fucosylation provides a host-derived glycan niche that supports beneficial commensals, SCFA-producing bacteria, epithelial barrier integrity, and controlled mucosal inflammation. In PWH on ART, senescence-associated induction of host FUCA1 and increased α-L-fucosidase activity promote degradation of intestinal α1,2-fucose despite preserved fucose-production signaling. Loss of α1,2-fucose reduces host-derived glycan support for beneficial microbes, contributing to depletion of SCFA-producing bacteria, lower SCFA production, microbial dysbiosis, impaired barrier resilience, inflammation, and biological aging phenotypes. Replenishment with the α1,2-fucose donor 2′FL restores microbial SCFA production, improves epithelial resilience to stress-mediated disruption, and promotes epithelial homeostasis programs, identifying intestinal α1,2-fucosylation as a modifiable host-microbiome axis in treated HIV infection.

## DISCUSSION

Persistent disruption of intestinal homeostasis is a defining feature of chronic treated HIV infection, yet the host mechanisms that convert mucosal stress into durable microbial and epithelial dysfunction remain incompletely understood.^18,63–68^ Here, we identify intestinal α1,2-fucosylation as a host-derived ecological and epithelial control point that is disrupted in PWH despite ART-mediated viral suppression. This defect occurred despite a similar FUT2 secretor genotype distribution between PWH and PWoH, indicating that reduced intestinal α1,2-fucose in treated HIV is primarily acquired rather than genetically predetermined. Mechanistically, the defect was not explained by increased microbial fucosidase capacity. Instead, our data suggest a host senescence-associated fucose-degradation program as an underlying mechanism of this defect. Functionally, lower intestinal α1,2-fucose was associated with depletion of SCFA-producing bacteria, reduced microbial diversity, increased inflammatory and biological aging phenotypes, and impaired epithelial barrier-associated signatures. Finally, replenishing α1,2-fucose using the human-milk-oligosaccharide-derived donor 2′FL restored microbial SCFA production, improved organoid resilience to inflammatory and ethanol-mediated disruption, and shifted epithelial transcriptional programs toward reduced inflammatory stress and improved epithelial organization. Together, these findings position intestinal α1,2-fucosylation as a modifiable host-microbiome axis in treated HIV infection.

A central implication of this study is that dysbiosis in chronic disease should not be viewed solely as a microbe-autonomous process.^69–71^ The host epithelium can actively regulate microbial community structure and function by controlling the availability of glycans that serve as nutrient cues, colonization determinants, and signals of mucosal status.^69,72,73^ In this framework, α1,2-fucose is part of a host program that helps stabilize a beneficial microbial ecosystem.^23,24,26^ Prior work has established that epithelial fucosylation is induced by host-microbe immune crosstalk and supports commensal fitness while restraining pathogenic behavior.^24,26,28,74^ Our findings extend this concept to treated HIV infection and suggest that disruption of host glycan ecology may be an upstream contributor to persistent dysbiosis. Notably, depletion of α1,2-fucose was one of many glycomic alterations observed in intestinal tissues from PWH compared with controls (Figure 1), suggesting that treated HIV is associated with broad and functionally relevant remodeling of the intestinal glycome. Defining how individual glycomic alterations affect epithelial integrity, microbial ecology, and intestinal homeostasis during treated HIV infection warrants further investigation. More broadly, these data support a model in which chronic mucosal stress can be translated into long-lasting microbial dysfunction through depletion of host-derived ecological support.

Our findings also argue that the α1,2-fucose defect in treated HIV is driven more by enhanced degradation than by a simple collapse of the fucosylation machinery. Reduced mucosal fucosylation could, in principle, result from impaired biosynthesis or increased removal of terminal fucose residues.^75,76^ However, microbial α-fucosidase capacity was not increased in PWH on ART, and genes involved in IL-22 signaling, GDP-fucose synthesis, and fucose transport were preserved or increased. These observations argue against a model in which reduced α1,2-fucose is caused primarily by loss of microbial fucosidase activity or global failure of epithelial fucose-production pathways. Instead, spatial transcriptomics demonstrated increased host FUCA1 expression within senescence-enriched intestinal niches, and induced epithelial senescence was sufficient to increase active α-L-fucosidase activity. Thus, senescence may not merely mark damaged mucosa; it may function as an effector program that actively remodels the glycan landscape encountered by gut microbes.^47,48,51^ Future studies should define the upstream drivers of this senescence-associated fucose-degrading state, including the potential roles of HIV-mediated mucosal injury, persistent low-level viral or immune activation, ART-associated toxicity, diet, microbial metabolites, sexual practices, and other demographic or environmental factors.

This catabolic model has broader conceptual significance. Chronic inflammatory and aging-associated diseases are frequently accompanied by epithelial stress, barrier dysfunction, and altered microbial ecology, but the molecular interfaces linking these processes remain poorly defined.^77–80^ Our data suggest that host glycan remodeling is one such interface. Although demonstrated here in treated HIV, the underlying logic of this pathway, stress-induced induction of glycan-degrading programs, depletion of host-beneficial microbial substrates, and downstream loss of microbial metabolic support, may extend to other chronic mucosal disorders characterized by persistent epithelial stress, inflammaging, and barrier failure. This broader relevance warrants further investigation. Defining whether similar acquired losses of α1,2-fucosylation occur in inflammatory bowel disease, aging, metabolic disease, alcohol use disorder, circadian disruption, and other chronic inflammatory conditions may reveal whether mucosal defucosylation is a generalizable mechanism of epithelial-microbial dysregulation.

The direct epithelial consequences of fucose loss also warrant deeper investigation. Our study focused primarily on the host-microbiome consequences of α1,2-fucose depletion, especially loss of SCFA-producing bacteria and reduced microbial metabolic output. However, emerging evidence indicates that epithelial fucosylation can also protect intestinal tissue through microbiome-independent or partially microbiome-independent mechanisms.^81^ FUT2-dependent α1,2-fucosylation protects intestinal stem cells against inflammatory injury by restraining ER stress, oxidative stress, mitochondrial dysfunction, and apoptosis. These observations raise the possibility that intestinal α1,2-fucose depletion may impair tissue homeostasis through two interconnected routes: indirectly, by depleting microbial substrates required for SCFA-producing bacteria, and directly, by weakening epithelial stress tolerance, stem-cell fitness, glycoprotein quality control, and repair programs. Distinguishing microbiome-dependent from microbiome-independent consequences of mucosal fucose depletion in PWH should be investigated.

Lower intestinal α1,2-fucose tracked with depletion of multiple SCFA-producing taxa, reduced microbiome diversity, enrichment of putatively pro-inflammatory bacteria, and reduced expression of tight junction proteins associated with epithelial barrier integrity. These relationships are biologically coherent because SCFAs, particularly butyrate, support colonocyte energetics, epithelial renewal, tight junction maintenance, and anti-inflammatory signaling. Thus, α1,2-fucose loss is well positioned to contribute to a feed-forward loop in which epithelial stress depletes a host-derived nutrient cue, beneficial fermenters decline, SCFA output falls, and barrier fragility and inflammation are further amplified. This model helps reconcile the long-standing question of whether dysbiosis in HIV is primarily a cause or consequence of mucosal injury. Our data suggest that both may be true: epithelial stress may initiate ecological disruption through glycan loss, and the resulting microbial metabolic defect may then reinforce epithelial dysfunction.

The associations with inflammatory and biological aging phenotypes place this pathway within a clinically meaningful network of host dysfunction. Lower α1,2-fucose levels were linked to markers of microbial translocation, epithelial injury, myeloid activation, systemic biological age acceleration, and intestinal biological aging. These correlations do not establish causality, but they support the idea that the intestinal α1,2-fucose defect is not an isolated mucosal abnormality. Rather, it appears embedded within a broader state of inflammaging and reduced tissue resilience. One plausible interpretation is that loss of α1,2-fucose contributes to a mucosal environment permissive for chronic immune stimulation and age-accelerating pathology. An alternative, not mutually exclusive, interpretation is that biological aging and mucosal senescence intensify fucose depletion, creating a self-reinforcing circuit linking epithelial aging, microbial dysfunction, and inflammation. This possibility is especially compelling in treated HIV, where premature aging signatures persist despite virologic control.

The rescue experiments strengthen the mechanistic and translational relevance of this model. Stool from PWH produced lower levels of SCFAs during anaerobic fermentation, yet retained the capacity to respond to α1,2-fucose replenishment in the form of 2′FL. This restoration of microbial metabolic output was accompanied by improved resilience of colon organoids exposed to inflammatory or ethanol-mediated stress, indicating recovery of barrier-supportive function. 2′FL has a favorable safety profile, is naturally abundant in human milk, and is already used in nutritional formulations, making it a plausible candidate for repurposing as a microbiome-directed glycan intervention. Future studies should determine whether 2′FL or other strategies that restore intestinal α1,2-fucose availability can improve microbial metabolic output, barrier function, inflammatory tone, and aging-related phenotypes *in vivo*.

### Limitations of the study

First, the human analyses are largely cross-sectional and cannot establish temporal directionality. Second, while our data strongly implicate senescence-associated FUCA1 and α-L-fucosidase activity in the loss of intestinal α1,2-fucose, additional perturbation studies will be required to define necessity *in vivo*, including direct inhibition or genetic targeting of FUCA1, senescence pathways, or fucose-degrading activity. Third, the fermentation and organoid experiments model key components of the host-microbiome interface but cannot fully capture the complexity of the intact intestinal mucosa, including immune cells, vascular signals, mucus architecture, and luminal flow. Fourth, although our study incorporated matched controls and multi-compartmental sampling, residual confounding is difficult to eliminate in human microbiome studies, particularly in HIV. Finally, although our lectin microarray data identify reduced α1,2-fucose-associated glycan signatures in intestinal tissues and epithelial crypts, these analyses do not fully resolve the relative contributions of epithelial cell-surface glycans versus mucus-associated mucin glycans. Future mucus-focused glycomic and spatial analyses will be required to determine how α1,2-fucose loss affects the mucus niche and mucus-associated microbial ecology in PWH.

### Conclusions

Our findings are the first to identify acquired loss of intestinal α1,2-fucosylation as a feature among treated PWH and position this defect at the intersection of epithelial senescence, host glycan degradation, microbial metabolism, barrier integrity, inflammation, and biological aging. More broadly, they suggest that host glycan ecology is a fundamental layer of host control over mucosal symbiosis and that its disruption may represent a mechanism by which chronic epithelial stress is converted into durable host-microbiome dysfunction. By demonstrating that this axis can be functionally rescued through glycan replenishment, our study establishes intestinal glycan remodeling not only as a disease mechanism among PWH but also as a tractable framework for restoring host-microbiome homeostasis among persons with chronic mucosal disease.

## METHODS

### Ethics Statement

Ileal and colonic biopsies, blood, and stool samples were collected from 25 PWH receiving ART and 22 PWoH of similar age, sex, body mass index (BMI), and race/ethnicity (**Table S1**) at Rush University Medical Center. The study protocol was approved by the Institutional Review Board at Rush University Medical Center (ORA# 19020710), and all participants provided written informed consent. Participants completed a detailed questionnaire that captured demographic information, and medical history. Samples were excluded from individuals reporting specialized diets, including vegan, vegetarian, gluten-free, Paleo, or specific carbohydrate diets, because these dietary patterns can influence gut microbial composition. Samples were also excluded from individuals with celiac disease.

### Patient and Public Involvement

The public were not involved in the design, or conduct, or reporting, or dissemination plans of our research.

### FUT2 secretor genotype analysis

Genomic DNA was isolated from PBMCs using the AllPrep DNA/RNA/miRNA Kit (Qiagen, catalog# 80224) according to the manufacturer’s instructions. FUT2 secretor status was inferred from the FUT2 rs516246 genotype, as previously described.^82^ Individuals with the TT genotype were classified as non-secretors, whereas those with CC or CT genotypes were classified as secretors. Genotyping was performed using 10 ng of genomic DNA, a TaqMan SNP Genotyping Assay for rs516246 (Thermo Fisher Scientific, catalog# 4351379, assay ID C_2885835_10), and TaqMan Genotyping Master Mix (Thermo Fisher Scientific, catalog #4371355) on a QuantStudio 6 Real-Time PCR System. Cycling conditions were as follows: 60°C for 30 s, 95°C for 10 min, followed by 40 cycles of 95°C for 15 s and 60°C for 1 min.

### Lectin microarray-based glycomic profiling

Glycomic profiling was performed using flash-frozen biopsy tissue samples as well as isolated crypts. Samples were washed with 500 μL of phosphate-buffered saline (PBS) containing 10 μL/mL protease inhibitor cocktail (Thermo Fisher Scientific, catalog# 87786). Samples were centrifuged at 800 xg for 5 min at 4°C, the supernatant was removed, and the wash step was repeated three additional times. After the final wash, pellets were resuspended in 300 μL of PBS containing protease inhibitor and homogenized using a pellet grinder (Sigma-Aldrich, catalog# Z359971-1EA). Homogenates were centrifuged at 1,500 xg for 5 min at 4°C, and the supernatant was discarded. Pellets were then resuspended in 300 μL of PBS containing 1% Triton X-100 (Thermo Fisher Scientific, catalog# A16046.AE) and protease inhibitor, followed by sonication at high intensity for 1 min using a Bioruptor Plus sonication device (Diagenode, catalog# B01020014). After sonication, samples were centrifuged at 12,000 xg for 10 min at 4°C, and protein-containing supernatants were collected, aliquoted, and stored at −80°C until analysis. Protein concentrations were determined using the Micro BCA Protein Assay Kit (Thermo Fisher Scientific, catalog# 23235) according to the manufacturer’s instructions. After quantification, 0.4 μg of protein was aliquoted into low-protein-binding tubes and labeled with 40 μg of Cy3 dye (Cytiva, catalog #PA23001) by incubation for 1 h at room temperature. Excess Cy3 dye was removed using 7K MWCO Zeba Spin Desalting Columns. Columns were equilibrated by washing three times with 300 μL of Tris-buffered saline (TBS), followed by centrifugation at 1,500 xg for 1 min at 4°C. Labeled samples were then applied to the columns and centrifuged at 1,500 xg for 2 min. After cleanup, samples were incubated for an additional 1 h, and the concentration was adjusted to 0.5 μg/mL in probing solution consisting of 25 mM Tris-HCl, pH 7.5, 140 mM NaCl, 2.7 mM KCl, 1 mM CaCl, 1 mM MnCl, and 1% Triton X-100. Labeled protein samples were hybridized overnight to lectin microarray chips containing 96 lectins (**Table S2**). Chips were scanned for fluorescence intensity using a Bio-Rex Scan 200 fluorescence scanner. Lectin-binding signals were analyzed using global normalization.

### Stool shotgun metagenomic sequencing and microbial α-fucosidase analysis

Metagenomic DNA extracted from fecal samples was sequenced on the Illumina NovaSeq 6000 platform, yielding paired-end reads. Standard demultiplexing and initial quality assessment were performed by the default workflows in Sunbeam.^83^ Raw reads were processed to remove Nextera-XT adapter sequences and low-quality bases using Trimmomatic (v0.33) with a sliding window quality threshold. Quality profiles before and after trimming were monitored using FastQC. Low complexity reads were removed using a customized script in Sunbeam. High-quality reads were subsequently aligned to the human reference genome (hg38) and the PhiX genome using BWA, and matching reads were removed to eliminate host-derived and sequencing control contamination. A reference collection of bacterial proteins annotated as α-L-fucosidase across taxa was downloaded from NCBI RefSeq on May 10, 2024. Metagenomic shotgun reads were aligned to this protein database using BLASTX with thresholds of percent identity >80%, alignment length >30 amino acids, and e-value <1×10^-8^. A subset of alignments was manually inspected and independently verified using the NCBI BLASTX web server to confirm annotation specificity. For each sample, aligned reads were aggregated across paired-end reads and normalized by the total number of quality-filtered reads in the library to generate Reads Per Million (RPM) values, providing estimates of α-L-fucosidase abundance and allowing comparisons across samples.

### Spatial transcriptomic profiling of intestinal tissues

Formalin-fixed paraffin-embedded (FFPE) human gut tissue sections were processed for spatial transcriptomics using the 10x Genomics Visium CytAssist platform with Visium V4 FFPE v2 chemistry. Tissue sections were mounted on Visium V4 slides and hybridized with the Visium Human Transcriptome Probe Set v2.0 (GRCh38-2020-A). Probe hybridization, ligation, and library preparation were performed according to the manufacturer’s protocol for FFPE tissue. CytAssist imaging was used for tissue registration and spatial barcode transfer, and brightfield images were acquired for each section. Generated libraries were subsequently sequenced in NUSeq Core on the Illumina Novaseq 6000 platform. Sequencing quality was high, with Q30 scores exceeding 96% across barcodes, probe reads, and UMIs, and 100% valid UMIs.

Raw sequencing data were processed using Space Ranger (v3.0.0). For each sample, spaceranger count was run with the human reference GRCh38-2020-A genome, also providing the Visium Human Transcriptome Probe Set v2.0. CytAssist and brightfield images were provided for tissue registration. Visium spatial transcriptomics data from four human gut samples (two HIV-positive and two HIV-negative) were loaded to Seurat (v.5.1.047) R package.^84^ Spots with zero counts were removed, and each sample was normalized using SCTransform. Mitochondrial and ribosomal gene percentages were calculated as quality metrics. Samples were merged and integrated using Harmony^85^ to correct for sample-level batch effects, grouping by donor identity. Principal component analysis (PCA) was performed, followed by UMAP and t-SNE embeddings using the first 15 Harmony-corrected dimensions. Unsupervised clustering was performed with the Leiden algorithm at resolution 0.3. Cluster marker genes were identified using Seurat’s FindAllMarkers function with MAST as the statistical framework,^86^ controlling for donor as a latent variable. HIV status–associated differential expression was assessed using FindMarkers with MAST to generate ranked gene lists for downstream enrichment analysis.

Custom gene set enrichment was performed against a curated gene list organized by biological category and subpathway module. Module scores for gene sets of interest (UPR/ER chaperones, Epithelial stress/UPR, IL-22 signaling) were computed using Seurat’s AddModuleScore. Senescence scoring analysis was completed in Python 3.10.18, using SenePy version 1.0.1.^52^ The hub settings species= “human” and hubs.metadata.tissue= “intestine” were used as backgrounds for senescence scoring. The hubs.get_genes function was used to select the categories “intestine” and “epithelial cell” for scoring. Raw spatial transcriptomics data for all samples were concatenated into a single anndata object and preprocessed using standard scanpy functions (scanpy.preprocessing.normalize, scanpy.preprocessing.log1p, scanpy.preprocessing.pca, scanpy.preprocessing.neighbors, scanpy.tools.umap, scanpy.tools.leiden). Senescence scores were then acquired using the SenePy command senepy.score_hub and scores were added to the anndata.obs metadata for visualization.

The relationship between high senescence (score > 0.1) and FUCA1 positivity (expression > 1; extracted from the normalized SCT assay) was modeled using a binomial generalized linear model (GLM), with HIV status included as an interaction term and log-transformed spot UMI counts as a covariate to control for sequencing depth. Estimated marginal means and pairwise contrasts stratified by HIV status were computed using the emmeans (v.2.0.3) R package with odds ratio scaling. To assess the continuous association between HIV status and senescence score, a robust linear model was fit using MM-estimation (rlm, MASS R package v.7.3-65), which provides resistance to violations of Gaussian assumptions caused by heavy-tailed residual distributions. The model included HIV status as the primary predictor and log-transformed normalized UMI counts as a covariate. Pairwise contrasts by HIV status were computed using lsmeans from emmeans.

### Caco-2 cell culture and chemically induced senescence

Caco-2 cells were cultured in Dulbecco’s modified Eagle medium (DMEM; Corning, catalog# MT10027CV) supplemented with 20% fetal bovine serum (FBS; Thermo Fisher Scientific, catalog# A5669701) and 1% penicillin-streptomycin-amphotericin B solution (Cytiva HyClone, catalog# SV3007901). Cells were passaged when they reached 80 to 90% confluence. For experiments, cells were detached using TrypLE Express Enzyme (Thermo Fisher Scientific, catalog# 1260401), and 0.1 × 10^6^ cells were seeded into each well of a 12-well plate. Cells were cultured for 3 days to allow attachment and recovery. Medium was then replaced, and cells were treated with 10 μg/mL bleomycin (Cayman Chemical, catalog# 13877) or 2.5 μM AZD1152-HQPA (Cayman Chemical, catalog# 11602) for 3 days to induce senescence. After treatment, culture supernatants were collected and stored at −80°C until analysis. Cells were detached with TrypLE and lysed in buffer containing 20 mM Tris-HCl, pH 7.4, 0.2% Triton X-100, and protease inhibitor. α-L-fucosidase activity was measured as previously described,^47^ with minor modifications. Briefly, 5 μL of culture supernatant or cell lysate was incubated with 45 μL of reaction buffer containing 0.2 M sodium citrate, pH 4.5, and 1 mM 4-methylumbelliferyl-α-L-fucopyranoside (Cayman Chemical, catalog# 39552) for 1 h at 37°C. To confirm assay specificity, parallel reactions were performed in the presence of an α-L-fucosidase inhibitor at 30mM of 1-Deoxyfuconojirimycin HCl (Cayman, catalog# MD06117). Reactions were stopped by adding 0.2 mL of 0.2 M sodium carbonate, followed by centrifugation at 2,000 xg for 10 min. Cleared supernatants were used to quantify 4-methylumbelliferone fluorescence at 355 nm excitation and 460 nm emission. 4-Methylumbelliferone concentrations were interpolated from a standard curve and normalized to the protein concentration in the corresponding sample. Senescence-associated β-galactosidase activity was measured by flow cytometry after 3 days of treatment using the CellEvent Senescence Green Flow Cytometry Assay Kit according to the manufacturer’s instructions (Thermo Fisher Scientific, catalog# C10841). Cells were acquired on a FACSymphony A5 Spectral Analyzer. Senescence-associated β-galactosidase activity was also visualized using the Senescence β-Galactosidase Staining Kit according to the manufacturer’s instructions (Cell Signaling Technology, catalog# 9860). To measure cell-surface uPAR expression, cells were detached with TrypLE after treatment, washed with PBS, and incubated for 30 min at 4°C with fluorophore-conjugated anti-CD87/uPAR monoclonal antibody (Thermo Fisher Scientific, catalog# 17-3879-42). Cells were then washed to remove unbound antibody and fixed with paraformaldehyde (BioLegend, catalog# 420801). Data were acquired on a FACSymphony A5 Spectral Analyzer.

### Measurement of plasma markers of microbial translocation, epithelial injury, and inflammation

Plasma levels were measured using electrochemiluminescence-based multiplex immunoassays from Meso Scale Discovery according to the manufacturer’s instructions. Assays included the U-PLEX Custom Biomarker Group 1 Human Assay (Meso Scale Discovery, catalog #K151AEM-2), U-PLEX Custom Immuno-Oncology Group 1 Human Assay (Meso Scale Discovery, catalog# K15067L-2), and Human MIG/CXCL9 Antibody Set (Meso Scale Discovery, catalog# F210I-3). Plasma concentrations of soluble CD14, soluble CD163, lipopolysaccharide-binding protein (LBP), and fatty acid-binding protein 2/intestinal fatty acid-binding protein (FABP2/I-FABP) were quantified using DuoSet ELISA kits according to the manufacturer’s instructions (R&D Systems, catalog# DY383-05, DY1607-05, DY870-05, and DFBP20, respectively).

### DNA methylation profiling and epigenetic clock estimation

For each sample, 300 ng of genomic DNA was subjected to bisulfite conversion using the EZ DNA Methylation Kit (Zymo Research) according to the manufacturer’s protocol. Bisulfite-converted DNA samples were randomized across wells of the Infinium HumanMethylationEPIC v1.0 BeadChip array. Samples were then amplified, hybridized to the array, stained, washed, and scanned using the Illumina iScan SQ system to generate raw image intensity files. Raw IDAT files from the MethylationEPIC array were imported into R and processed using the SeSAMe R package.^87^ DNA methylation beta values were then used to estimate multiple epigenetic aging and aging-related biomarkers, including Horvath’s multi-tissue DNAmAge clock,^61^ Horvath’s skin-and-blood clock,^88^ Levine’s DNAmPhenoAge clock,^89^ the Hannum DNAmAge clock,^90^ Lu’s DNA methylation-based telomere length estimator,^91^ and GrimAge.^92^ Principal component-based epigenetic clock estimates were calculated using the R script provided by Higgins-Chen et al., with 78,464 CpGs included for each sample in the beta-value matrix. Missing values required for clock calculation were imputed using mean imputation. DunedinPACE was calculated using the publicly available implementation provided by the developers.^93^

### Telomere length quantification

Telomere length in PBMCs was measured using the high-throughput quantitative fluorescence in situ hybridization (HT Q-FISH)-based Telomere Analysis Technology (TAT) platform developed by Life Length Technologies.^94^ Briefly, PBMCs were fixed and hybridized with a fluorescent peptide nucleic acid (PNA) probe complementary to telomeric repeat sequences. Following hybridization, cells were washed to remove nonspecific probe binding, and nuclei were counterstained with DAPI. Nuclear and telomeric fluorescence images were acquired using a high-content imaging system (Opera Phenix, PerkinElmer). For each field, maximum-intensity projection images were generated from multiple Z-stack images to improve telomere signal detection and quantification. Telomere fluorescence intensities were converted to estimated telomere length in base pairs using a standard regression curve generated from control cell lines with known telomere lengths. Data were processed using the proprietary TAT Analyzer software to generate telomere-associated variables (TAVs), including the percentage of telomeres within defined length ranges and the percentage of cells harboring telomeres within specified length ranges, and dispersion metrics for each sample.

### 16S rRNA sequencing and microbiome analysis of fecal, colonic, and ileal samples

DNA was extracted from approximately 200 mg of stool or available biopsy tissue using the DNeasy PowerSoil Pro Kit (Qiagen) and quantified using the Quant-iT PicoGreen Assay Kit. Barcoded primers targeting the V1-V2 region of the bacterial 16S rRNA gene were used to generate amplicon libraries. PCR reactions were performed in duplicate using Q5 High-Fidelity DNA Polymerase (New England Biolabs). For high microbial biomass samples, including fecal samples, each 50 μL PCR reaction contained 0.5 μM of each primer, 0.34 U Q5 polymerase, 1x reaction buffer, 0.2 mM dNTPs, and 5 μL of template DNA. Cycling conditions were as follows: 98°C for 1 min; 20 cycles of 98°C for 10 s, 56°C for 20 s, and 72°C for 20 s; followed by a final extension at 72°C for 8 min. For low microbial biomass samples, including colonic and ileal biopsy tissues, PCR reactions were performed using the same conditions except that 10 μL of template DNA and 25 amplification cycles were used. Duplicate PCR products from each sample were pooled and purified using a 1:1 volume of SPRI beads. Purified amplicons were quantified using PicoGreen and pooled in equimolar amounts. The final pooled library was sequenced on an Illumina MiSeq platform using 2 x 250 bp paired-end chemistry. Extraction blanks and DNA-free water controls were processed through the same amplification and purification workflow to assess environmental and reagent-derived contamination. Positive controls consisting of eight synthetic 16S rRNA gene fragments, generated as gene blocks and combined at known abundances, were included to evaluate amplification and sequencing performance. No appreciable contamination was observed in negative controls, and positive control samples generated the expected read distributions.

Sequence data were processed using QIIME 2 version 2019.7.^95^ Paired-end reads were denoised and resolved into amplicon sequence variants using DADA2, with both forward and reverse reads truncated at 240 nucleotides.^96^ Taxonomic assignments were generated against the SILVA reference database version 132^97^ using the naïve Bayes classifier implemented in scikit-bio with default parameters.^98^ Numbered taxa, such as Prevotella 2, represent distinct taxonomic groups as defined in the SILVA taxonomy. QIIME output files were imported into R for downstream statistical analysis.^99^ To evaluate microbial groups relevant to intestinal barrier function and inflammation, genus-level 16S rRNA relative abundance data were used to generate literature-curated bacterial guilds. Taxa were classified based on reported involvement in SCFA production, butyrate production, or putative pro-inflammatory activity (**Table S3**).^100–106^

### Immunofluorescence assessment of tight junction proteins in ileal and colonic tissues

Ileal and colonic biopsies were embedded in optimal cutting temperature (OCT) compound and sectioned at 5 μm thickness. Tissue sections were fixed in a 1:1 acetone/methanol solution at −20°C for 2 min, air-dried, and rehydrated in 1x PBS for 10 min. Sections were then permeabilized with 0.2% Triton X-100 in PBS for 5 min, washed, and blocked for 1 h at room temperature using 2% non-fat dry milk in PBS. Sections were incubated for 1 h at 37°C with primary antibodies against ZO-1 (Invitrogen, catalog #61-7300) or occludin (Invitrogen, catalog# 33-1500) diluted in 2% non-fat dry milk. After washing, sections were incubated for 1 h at 37°C with Alexa Fluor 488-conjugated donkey anti-rabbit IgG (Invitrogen, catalog# A-21206) or Alexa Fluor 488-conjugated donkey anti-mouse IgG (Invitrogen, catalog# A-21202) secondary antibodies diluted 1:250. Sections were then washed, counterstained with DAPI for 3 min, washed again, and mounted using Fluoromount Aqueous Mounting Medium (Sigma, catalog# F4680). Images were acquired at 40× magnification using a Zeiss Axio Observer 7 digital deconvolution immunofluorescence microscope. For each sample, at least five stained tissue fields were imaged and used to quantify the relative abundance of each tight junction marker and to select representative images. Immunofluorescence images were evaluated by two independent observers who were blinded to participant group assignment.

### Anaerobic stool fermentation with 2′-FL

*In vitro* fecal fermentations were performed following a previously described methodology with modifications.^107^ Carbonate-phosphate buffer was prepared, sterilized by autoclaving (121°C for 20 min), reduced with cysteine hydrochloride (0.25 g/L), and placed in an anaerobic chamber overnight prior to fermentation. On the day of the experiment, fecal samples were homogenized with carbonate-phosphate buffer at a 1:3 (wt/vol) ratio and filtered through four layers of cheesecloth. Fecal inocula were prepared individually for each donor. Fermentations were conducted in Balch tubes containing 50 mg of substrate and 4 mL of carbonate-phosphate buffer. Fructooligosaccharides (FOS; Sigma-Aldrich, Catalog# F8052) and 2′FL (Layer Origin Nutrition, Catalog# HMO 2’FL) were used as fermentation substrates, while tubes without added substrate served as blank controls. Subsequently, 1 mL of fecal inoculum was added to each tube. Tubes were sealed with butyl rubber stoppers and aluminum crimps and incubated anaerobically at 37°C with shaking (150 rpm) for 48 h. Baseline and post-fermentation aliquots were collected and stored at −80°C until further use and analysis. All sample handling was performed under anaerobic conditions (85% N, 10% H, and 5% CO).

### SCFA analysis of fermentation supernatants

Fermentation supernatants were prepared as previously described^107^ and analyzed for SCFA content using a gas chromatograph equipped with a flame ionization detector (GC-FID 7890A; Agilent Technologies) and a fused-silica capillary column (Nukol, Supelco # 40369-03A). The operating conditions were as follows: injector temperature, 230°C; initial oven temperature, 100°C; temperature ramp, 8°C/min to 200°C; and a final hold of 3 min at 200°C. Helium was used as the carrier gas at a flow rate of 0.75 mL/min. SCFAs were quantified using external calibration standards for acetate (A38S), propionate (A258), and butyrate (AC108111000), with 4-methylvaleric acid (AAA1540506) used as the internal standard (Fisher Scientific). Quantification was based on peak area ratios relative to the internal standard.

### Human intestinal organoid culture

Sigmoid colon–derived organoids (colonoids) were generated by isolating intestinal stem cells from a healthy control subject. Cells were embedded in Matrigel and supplemented with defined growth factors to promote organoid formation, as previously described.^9,108,109^ For this study, organoid polarity was reversed using an established method to produce apical-out organoids, enabling direct access to the apical epithelium while maintaining three-dimensional structure.^110^ Apical-out organoids were pretreated for 24 hours (1:100 dilution) with stool fermentation supernatants. These supernatants were obtained either from PWoH or PWH on ART, generated from 48-hour blank fermentations or fermentations conducted in the presence of 2′FL substrate. Following pretreatment, organoids were exposed for 3 hours to either a cytokine cocktail containing IFN-γ and TNF-α (each at 6 ng/mL) or ethanol (0.2%). Organoids were subsequently pelleted, washed twice with 1× PBS, and resuspended in growth medium containing 4 kDa FITC-dextran (2 mg/mL). Live imaging was performed using differential interference contrast (DIC) and fluorescence microscopy with a 20x objective on a Zeiss Axio Observer 7 digital deconvolution immunofluorescence microscope (Carl Zeiss) using Zen Blue 3.10 software. FITC-dextran fluorescence intensity was quantified as a measure of epithelial barrier function, with increased fluorescence indicating enhanced permeability and tight junction disruption.^9^

### RNA-seq of intestinal organoids exposed to fermentation supernatants

Total RNA was extracted from intestinal organoids using the Quick-DNA/RNA Miniprep Plus Kit according to the manufacturer’s instructions (Zymo, catalog# D7003T). RNA concentration and purity were assessed using a Qubit Fluorometer, and RNA integrity was evaluated using Agilent Bioanalyzer 2100, with samples meeting the required quality threshold selected for sequencing. RNA-seq libraries were prepared from 100ng of total RNA using Illumina Stranded mRNA Seq Library Prep kit, with polyA capture according to the manufacturer’s protocol. Indexed libraries were quantified, pooled at equimolar concentrations, and sequenced on an Illumina platform using NovaSeq X Plus sequencing to generate raw FASTQ files for downstream bioinformatic analysis.

Raw RNA-seq data were processed using the RNA-seq analysis pipeline implemented in TWINE RNA Suite v1.0. Raw read quality was assessed using FastQC v0.12.1, followed by adapter and quality trimming with fastp v0.23.2. Adapters were auto-detected, poly-G artifacts were removed, low-quality reads were filtered, 3′ bases with Phred quality scores <30 were trimmed, and reads shorter than 20 bp after trimming were excluded. Cleaned reads were aligned to the human GRCh38 reference genome from Gencode using the splice-aware aligner STAR v2.7.10b within the STAR-Salmon workflow. STAR was run with transcriptome BAM and gene-count output generation enabled to support downstream transcript-level quantification. Transcript abundance was quantified using Salmon v1.11.4, with correction for sample-specific sequence and GC-content biases. Salmon outputs were imported into DESeq2 v1.42.1 using tximport for gene-level differential-expression analysis.

Differential expression was performed using DESeq2, which models RNA-seq count data with a negative binomial framework. Genes were retained for analysis if they had at least five reads in at least 30% of samples. Counts were normalized using the median-of-ratios method to account for differences in sequencing depth and RNA composition. Differential expression between prespecified paired comparisons was assessed using Wald tests, and P values were adjusted for multiple testing using the Benjamini–Hochberg false-discovery-rate procedure.

### Statistical analysis

Statistical analyses were performed using GraphPad Prism version 10, R, Python, and bioinformatic packages described above, as appropriate for each dataset. For standard group comparisons, normality was assessed using the Shapiro–Wilk test when parametric analyses were considered. Comparisons between PWH on ART and PWoH were performed using two-sided Mann–Whitney U tests for nonparametric continuous variables and Fisher’s exact test for categorical variables, including FUT2 secretor status. Paired comparisons, including SCFA production in matched stool fermentations performed with or without 2′FL, were analyzed using Wilcoxon matched-pairs signed-rank tests. For experiments involving more than two groups, including chemically induced senescence assays in Caco-2 cells and organoid permeability analyses, one-way ANOVA or mixed-effects analyses were used as indicated in the corresponding figure legends, with multiple comparisons corrected using the two-stage step-up method of Benjamini, Krieger, and Yekutieli. Correlations between intestinal α1,2-fucose levels, inflammatory markers, biological aging measures, microbiome features, SCFA-producing bacterial groups, microbial diversity metrics, tight junction markers, and fermentation outputs were assessed using Spearman’s rank correlation.

For spatial transcriptomic analyses, differential expression was assessed using Seurat/MAST with donor identity included as a latent variable, and P values were adjusted using the Benjamini–Hochberg false-discovery-rate procedure. The association between high senescence score and FUCA1 positivity was modeled using a binomial generalized linear model with HIV status as an interaction term and log-transformed spot UMI counts included as a covariate; estimated marginal means and pairwise contrasts were computed using the emmeans R package. Continuous senescence-score differences by HIV status were assessed using robust linear models with MM-estimation, with normalized UMI counts included as a covariate. For microbiome and shotgun metagenomic analyses, group-level differential abundance comparisons were performed using two-sided Mann-Whitney U tests, with Benjamini-Hochberg correction applied where indicated. For organoid RNA-seq, differential expression was performed using DESeq2, with genes retained if they had at least five reads in at least 30% of samples; normalized counts were analyzed using Wald tests, and P values were adjusted using the Benjamini–Hochberg false-discovery-rate procedure. Exact n values, the definition of n, precision measures, and the statistical test used for each panel are provided in the corresponding figure legends. No formal sample-size estimation was performed.

## ACKNOWLEDGMENTS

We would like to thank study participants. This study is supported by the National Institutes of Health (NIH) R01DK123733 to M.A-M, A.K, and A.L as well as R01AA031197 to A.K and M-A-M. M.A-M is also supported by NIH grants (R01AI165079, R01AI189353, and R01AG092241). A.K is also supported by NIAAA (R24AA026801). M.A-M is a member of the NIH-funded BEAT-HIV Martin Delaney Collaboratory to cure HIV-1 infection (1UM1Al126620). This research was also supported in part by philanthropic funding from Mr. and Mrs. Larry Field, Mr. and Mrs. Glass, Mrs. Marcia and Mr. Silas Keehn, the Sklar Family, the Johnson Family, and Mr. Harlan Berk to A.K.

## AUTHOR CONTRIBUTIONS

M.A.-M. and A.K. conceived and designed the study. L.B.G., M.W.S., L.Z, N.C, S.S., and P.A.E. performed the experiments. Sh.Sh., A.L.L., F.J.P., and M.V. selected the clinical samples and contributed to clinical data interpretation. M.J.C. supervised the epigenetic aging clock analyses. H.T contributed to the lectin array data interpretation. T.M.C.J. and B.H. performed the fermentation assays. N.A. analyzed the stool shotgun metagenomic data. R.L.R., J.M.H, E.Z, and T.J.H performed and analyzed the spatial transcriptomic analyses. L.B.G. and M.A.-M. had full access to all data and wrote the first draft of the manuscript. All authors read and approved the final article and take responsibility for its content.

