## Supplementary Materials for "Senescence-associated loss of intestinal α1,2-fucose disrupts a modifiable host-microbiome homeostasis axis in people with HIV"

**Table S1. Demographic and clinical characteristics of study participants; related to Figure 1.**

| Characteristic | PWoH (n=22) | PWH on ART (n=25) |
|---|---|---|
| Age, years, median (IQR) | 53 (11) | 54 (12) |
| BMI, kg/m², median (IQR) | 28.01 (4.22) | 28 (9.9) |
| **Race, n (%)** | | |
| African American | 8 (36.4%) | 9 (36%) |
| White | 11 (50%) | 13 (52%) |
| Other | 3 (13.6%) | 3 (12%) |
| **Ethnicity, n (%)** | | |
| Hispanic | 3 (13.6%) | 3 (12%) |
| Non-Hispanic | 19 (86.4%) | 22 (88%) |
| **Sex at birth, n (%)** | | |
| Male | 16 (72.7%) | 21 (84%) |
| Female | 6 (27.3%) | 4 (16%) |
| **Sexual orientation/behavior, n (%)** | | |
| Heterosexual | 20 (90.9%) | 7 (28%) |
| Homosexual | 0 (0%) | 13 (52%) |
| MSM | 0 (0%) | 5 (20%) |
| Unknown | 2 (9.1%) | 0 (0%) |
| **HIV-related characteristics** | | |
| CD4 count at sampling, cells/μL, median (IQR) | N/A | 593 (389) |
| Nadir CD4 count, cells/μL, median (IQR) | N/A | 216 (200) |
| Year of HIV diagnosis, median (IQR) | N/A | 2001 (13.5) |
| **Viral load at sampling, copies/mL, n (%)** | | |
| <20 | N/A | 15 (60%) |
| <40 | N/A | 7 (28%) |
| 41 | N/A | 1 (4%) |
| Not detected | N/A | 2 (8%) |
| **ART regimen at sampling, n (%)** | | |
| Abacavir, dolutegravir, and lamivudine | N/A | 3 (12%) |
| Bictegravir, emtricitabine, and tenofovir alafenamide | N/A | 11 (44%) |
| Dolutegravir and doravirine | N/A | 1 (4%) |
| Dolutegravir, emtricitabine, tenofovir alafenamide, and doravirine | N/A | 1 (4%) |
| Dolutegravir and lamivudine | N/A | 2 (8%) |
| Elvitegravir, cobicistat, emtricitabine, and tenofovir | N/A | 2 (8%) |
| Emtricitabine, rilpivirine, and tenofovir alafenamide | N/A | 4 (16%) |
| Dolutegravir, darunavir, cobicistat | N/A | 1 (4%) |
| **Diabetes, n (%)** | | |
| No | 22 (100%) | 21 (87.5%) |
| Yes | 0 (0%) | 2 (12.5%) |
| **Liver enzymes, U/L, median (IQR)** | | |
| AST | 18.5 (5.5) | 24 (14) |
| ALT | 18.5 (10) | 23 (17) |
| **Gut disease, n (%)** | | |
| No | 17 (77.3%) | 14 (56%) |
| Yes | 5 (22.7%) | 11 (44%) |
| GERD | 5 (100%) | 8 (72.7%) |
| Crohn's disease | 0 (0%) | 1 (9.1%) |
| Other | 0 (0%) | 1 (9.1%) |
| Not available | 0 (0%) | 1 (9.1%) |
| **Heart disease, n (%)** | | |
| No | 15 (68.2%) | 5 (24%) |
| Yes | 7 (31.8%) | 19 (76%) |
| Hypertension | 3 (42.9%) | 9 (47.4%) |
| Hyperlipidemia | 1 (14.3%) | 2 (10.5%) |
| Other | 1 (14.3%) | 1 (5.3%) |
| More than one condition | 2 (28.6%) | 7 (36.8%) |
| **Neurological disease, n (%)** | | |
| No | 19 (86.4%) | 13 (52%) |
| Yes | 3 (13.6%) | 12 (48%) |
| ADD, epilepsy, stroke | 0 (0%) | 0 (0%) |
| Shingles | 0 (0%) | 3 (25%) |
| Bipolar disorder | 0 (0%) | 2 (16.7%) |
| Depression | 0 (0%) | 1 (8.3%) |
| More than one condition | 0 (0%) | 3 (25%) |
| Other (Bell's palsy, migraine, neurocardiogenic pre-syncope) | 0 (0%) | 3 (25%) |
| **Hepatitis status, n (%)** | | |
| No | 21 (95.5%) | 17 (68%) |
| Yes | 0 (0%) | 6 (24%) |
| Unknown | 1 (4.5%) | 2 (8%) |
| Hepatitis A | 0 (0%) | 1 (16.7%) |
| Hepatitis B | 0 (0%) | 2 (33.3%) |
| Hepatitis C | 0 (0%) | 3 (50%) |
| **Smoking status, n (%)** | | |
| Never | 18 (81.8%) | 14 (56%) |
| Former | 2 (9.1%) | 7 (28%) |
| Current | 2 (9.1%) | 4 (16%) |
| **Drug use, excluding alcohol, n (%)** | | |
| None | 19 (86.4%) | 17 (68%) |
| Marijuana | 3 (13.6%) | 8 (32%) |

**Table S2.** Lectins used in the lectin microarray experiments; related to Figure 1.

| | Name | Species | Origin | Glycan specificity* |
|---|---|---|---|---|
| 1 | LFA | *Limax flavus* | Natural | Sia |
| 2 | WGA | *Triticum vulgaris* | Natural | (GlcNAc)n, polySia |
| 3 | PVL | *Psathyrella velutina* | Natural | Sia, GlcNAc |
| 4 | MAL | *Maackia amurensis* | Natural | α2-3Sia |
| 5 | MAH | *Maackia amurensis* | Natural | α2-3Sia |
| 6 | ACG | *Agrocybe cylindracea* | Natural | α2-3Sia |
| 7 | rACG | *Agrocybe cylindracea* | *Recombinant* | α2-3Sia |
| 8 | rGal8N | *Homo sapiens* | *Recombinant* | α2-3Sia |
| 9 | SNA | *Sambucus nigra* | Natural | α2-6Sia |
| 10 | SSA | *Sambucus sieboldiana* | Natural | α2-6Sia |
| 11 | TJAI | *Trichosanthes japonica* | Natural | α2-6Sia |
| 12 | rPSL1a | *Polyporus squamosus* | *Recombinant* | α2-6Sia |
| 13 | PHAL | *Phaseolus vulgaris* | Natural | GlcNAcβ1-6Man (Tetraantenna) |
| 14 | DSA | *Datura stramonium* | Natural | GlcNAcβ1-6Man (Tetraantenna) |
| 15 | TxLcl | *Tulipa gesneriana* | Natural | Galactosylated N-glycans up to triantenna |
| 16 | ECA | *Erythrina cristagalli* | Natural | βGal |
| 17 | RCA120 | *Ricinus communis* | Natural | βGal |
| 18 | rGal7 | *Homo sapiens* | *Recombinant* | Type1 LacNAc, chondroitin polymer |
| 19 | rGal9N | *Homo sapiens* | *Recombinant* | GalNAcα1-4Gal (A), PolyLacNAc |
| 20 | rGal9C | *Homo sapiens* | *Recombinant* | PolyLacNAc, Branched LacNAc |
| 21 | rC14 | *Gallus gallus domesticus* | *Recombinant* | Branched LacNAc |
| 22 | rDiscoidin II | *Dictyostelium dicodeum* | *Recombinant* | LacNAc, Galβ1-3GalNAc (T), GalNAc (Tn) |
| 23 | BPL | *Bauhinia purpurea alba* | Natural | Galβ1-3GlcNAc(GalNAc), α/βGalNAc |
| 24 | rCGL2 | *Homo sapiens* | *Recombinant* | GalNAcα1-3Gal (A), PolyLacNAc |
| 25 | PHAE | *Phaseolus vulgaris* | Natural | bisecting GlcNAc |
| 26 | GSLII | *Griffonia simplicifolia* | Natural | GlcNAcβ1-4Man |
| 27 | rSRL | *Sclerotium rolfsii* | *Recombinant* | Galβ1-3GalNAc (T), GalNAcα (Tn) |
| 28 | UDA | *Urtica dioica* | Natural | (GlcNAc)n |
| 29 | PWM | *Phytolacca americana* | Natural | (GlcNAc)n |
| 30 | rF17AG | *Escherichia coli* | *Recombinant* | GlcNAc |
| 31 | rGRFT | *Griffithia sp.* | *Recombinant* | Man |
| 32 | NPA | *Narcissus pseudonarcissus* | Natural | Manα1-3Man |
| 33 | ConA | *Canavalia ensiformis* | Natural | M3, Manα1-2Manα1-3(Manα1-6)Man, GlcNAcβ1-2Manα1-3(Manα1-6)Man |
| 34 | GNA | *Galanthus nivalis* | Natural | Manα1-3Man, Manα1-6Man |
| 35 | HHL | *Hippeastrum hybrid* | Natural | Manα1-3Man, Manα1-6Man |
| 36 | ASA | *Allium sativum* | Natural | Galβ1-4GlcNAcβ1-2Man |
| 37 | DBAI | *Dioscorea batatas* | Natural | High-man |
| 38 | CCA | *Castanea crenata* | Natural | Galactosylated N-glycans up to triantenna |
| 39 | Heltuba | *Helianthus t uberosus* | Natural | Manα1-3Man |
| 40 | rHeltuba | *Helianthus tuberosus* | *Recombinant* | Manα1-3Man |
| 41 | ADA | *Allomyrina dichtoma* | Natural | α2-6Sia, Forssman, A, B |
| 42 | VVAII | *Vicia villosa* | Natural | Man, Agalacto |
| 43 | rOrysata | *Oryza sativa* | *Recombinant* | Manα1-3Man, Highman, biantenna |
| 44 | rPALa | *Phlebodium aureum* | *Recombinant* | Man5, biantenna |
| 45 | rBanana | *Musa acuminata* | *Recombinant* | Manα1-2Manα1-3(6)Man |
| 46 | rCalsepa | *Calystegia sepium* | *Recombinant* | Biantenna with bisecting GlcNAc |
| 47 | rRSL | *Ralstonia solanacearum* | *Recombinant* | αMan, α1-2Fuc (H), α1-3Fuc ($Le^x$), α1-4Fuc ($Le^a$) |
| 48 | rBC2LA | *Burkholderia cenocepacia* | *Recombinant* | αMan, High-man |
| 49 | AOL | *Aspergillus oryzae* | Natural | α1-2Fuc (H), α1-3Fuc (Lex), α1-3Fuc (Lea) |
| 50 | AAL | *Aleuria aurantia* | Natural | α1-6Fuc (Core), α1-2Fuc (H), α1-3Fuc ($Le^x$), α1-3Fuc ($Le^a$) |
| 51 | rAAL | *Aleuria aurantia* | *Recombinant* | α1-6Fuc (Core), α1-2Fuc (H), α1-3Fuc ($Le^x$), α1-3Fuc ($Le^a$) |
| 52 | rPAIIL | *Pseudomonas aeruginosa* | *Recombinant* | αMan, α1-2Fuc (H), α1-3Fuc ($Le^x$), α1-4Fuc ($Le^a$) |
| 53 | rRSIIL | *Ralstonia solanacearum* | *Recombinant* | α1-2Fuc (H), α1-3Fuc ($Le^x$), α1-3Fuc ($Le^a$) |
| 54 | rPTL | *Pholiota terrestris* | *Recombinant* | α1-6Fuc |
| 55 | PSA | *Pisum sativum* | Natural | α1-6Fuc up to biantenna |
| 56 | LCA | *Lens culinaris* | Natural | α1-6Fuc up to biantenna |
| 57 | rAOL | *Aspergillus oryzae* | *Recombinant* | α1-2Fuc (H), α1-3Fuc (Lex), α1-3Fuc (Lea) |
| 58 | rBC2LCN | *Burkholderia cenocepacia* | *Recombinant* | Fuc α1-2Galb1-3GlcNAc (GalNAc) |
| 59 | LTL | *Lotus tetragonolobus* | Natural | Fuc ($Le^x$, $Le^y$) |
| 60 | UEAI | *Ulex europaeus* | Natural | α1-2Fuc |
| 61 | TJAII | *Trichosanthes japonica* | Natural | α1-2Fuc |
| 62 | MCA | *Momordica charantia* | Natural | α1-2Fuc |
| 63 | GSLI | *Griffonia simplicifolia* | Natural | αGalNAc (A, Tn), αGal (B) |
| 64 | PTLI | *Psophocarpus tetragonolobus* | Natural | αGalNAc (A, Tn) |
| 65 | GSLIA4 | *Griffonia simplicifolia* | Natural | αGalNAc (A, Tn) |
| 66 | rGC2 | *Geodia cydonium* | *Recombinant* | α1-2Fuc (H), αGalNAc (A), αGal (B) |
| 67 | GSLIB4 | *Griffonia simplicifolia* | Natural | αGal (B) |
| 68 | rMOA | *Marasmius oreades* | *Recombinant* | αGal (B) |
| 69 | EEL | *Euonymus europaeus* | Natural | αGal (B) |
| 70 | rPAIL | *Pseudomonas aeruginosa* | *Recombinant* | α,βGal, αGalNAc (Tn) |
| 71 | LEL | *Lycopersicon esculentum* | Natural | Polylactosamine, (GlcNAc)n |
| 72 | STL | *Solanum tuberosum* | Natural | Polylactosamine, (GlcNAc)n |
| 73 | rGal3C | *Homo sapiens* | *Recombinant* | LacNAc, polylactosamine |
| 74 | rLSLN | *Laetiporus sulphureus* | *Recombinant* | LacNAc, polylactosamine |
| 75 | rCGL3 | *Coprinopsis cinerea* | *Recombinant* | LacDiNAc |
| 76 | PNA | *Arachis hypogaea* | Natural | Galβ1-3GalNAc (T), GalNAcα (Tn) |
| 77 | ACA | *Amaranthus caudatus* | Natural | Galβ1-3GalNAc (T), GalNAcα (Tn) |
| 78 | HEA | *Hericium erinaceum* | Natural | Galβ1-3GalNAc (T) |
| 79 | ABA | *Agarics bisporus* | Natural | Galβ1-3GalNAc (T), GlcNAc |
| 80 | Jacalin | *Artocarpus integrifolia* | Natural | Galβ1-3GalNAc (T), GalNAcα (Tn) |
| 81 | MPA | *Maclura pomifera* | Natural | Galβ1-3GalNAc (T), GalNAcα (Tn) |
| 82 | HPA | *Helix pomatia* | Natural | αGalNAc (A, Tn) |
| 83 | VVA | *Vicia villosa* | Natural | α,βGalNAc (A, Tn, LacDiNAc) |
| 84 | DBA | *Dolichos biflorus* | Natural | α,βGalNAc (A, Tn, LacDiNAc) |
| 85 | SBA | *Glycine max* | Natural | α,βGalNAc (A, Tn, LacDiNAc) |
| 86 | rPPL | *Pleurocybella porrigens* | *Recombinant* | α,βGalNAc (A, Tn, LacDiNAc) |
| 87 | rCNL | *Clitocybe nebularis* | *Recombinant* | α,βGalNAc (A, Tn, LacDiNAc) |
| 88 | rXCL | *Xerocomus chrysenteron* | *Recombinant* | Core1,3, agalacto N-glycan |
| 89 | VVA I | *Vicia villosa* | Natural | GalNAcβ1-3(4)Gal |
| 90 | WFA | *Wisteria floribunda* | Natural | Terminal GalNAc, LacDiNAc |
| 91 | rABA | *Agarics bisporus* | *Recombinant* | Galβ1-3GalNAc (T), GlcNAc |
| 92 | rDiscoidin I | *Dictyostelium Discodeum* | *Recombinant* | Gal |
| 93 | DBAIII | *Dioscorea batatas* | Natural | Maltose |
| 94 | rMalectin | *Homo sapiens* | *Recombinant* | Glcα1-2Glc |
| 95 | CSA | *Oncorhynchus keta* | Natural | Rhamnose, Galα1-4Gal |
| 96 | FLAG-EW29Ch-E20l | *Lumbricus terrestris* | *Recombinant* | 6-sulfo-Gal |

*Abbreviations: Gal (D-galactose), GalNAc (N-acetyl-galactosamine), GlcNAc (N-acetyl-glucosamine), Fuc (L-fucose), Glc (D-glucose), Sia (Sialic acid), LacNAc (N-acetyl-lactosamine).

**Table S3. Lists of bacteria used to calculate the relative abundance of (A) SCFA-producing bacteria, (B) butyrate-producing bacteria, and (C) pro-inflammatory bacteria; related to Figure 5.**

| *SCFA-producing bacteria (n = 85)* | *Butyrate-producing bacteria (n = 50)* | *Pro-inflammatory bacteria (n = 91)* |
|---|---|---|
| *p__Firmicutes g__Blautia* | *p__Firmicutes g__Faecalibacterium* | *p__Bacteroidetes g__Bacteroides* |
| *p__Firmicutes g__Faecalibacterium* | *p__Firmicutes g__[Ruminococcus] torques group* | *p__Firmicutes g__Streptococcus* |
| *p__Actinobacteria g__Collinsella* | *p__Firmicutes g__Subdoligranulum* | *p__Firmicutes g__Holdemanella* |
| *p__Firmicutes g__Subdoligranulum* | *p__Firmicutes g__[Ruminococcus] gnavus group* | *p__Fusobacteria g__Fusobacterium* |
| *p__Firmicutes g__[Eubacterium] hallii group* | *p__Firmicutes f__Lachnospiraceae* | *p__Bacteroidetes g__Parabacteroides* |
| *p__Firmicutes g__Dorea* | *p__Firmicutes g__Lachnoclostridium* | *p__Proteobacteria f__Enterobacteriaceae* |
| *p__Firmicutes g__Anaerostipes* | *p__Firmicutes g__Coprococcus 3* | *p__Proteobacteria g__Sutterella* |
| *p__Firmicutes g__[Ruminococcus] torques group* | *p__Firmicutes g__[Eubacterium] hallii group* | *p__Bacteroidetes g__Alistipes* |
| *p__Firmicutes g__Agathobacter* | *p__Firmicutes g__Agathobacter* | *p__Firmicutes g__Catenibacterium* |
| *p__Firmicutes g__Fusicatenibacter* | *p__Firmicutes g__Roseburia* | *p__Firmicutes g__Staphylococcus* |
| *p__Firmicutes g__[Ruminococcus] gauvreauii group* | *p__Firmicutes g__Anaerostipes* | *p__Proteobacteria g__Haemophilus* |
| *p__Firmicutes f__Lachnospiraceae* | *p__Firmicutes g__Coprococcus 1* | *p__Proteobacteria g__Escherichia-Shigella* |
| *p__Firmicutes g__Ruminococcus 2* | *p__Firmicutes g__Fusicatenibacter* | *p__Spirochaetes g__Brachyspira* |
| *p__Firmicutes g__Turicibacter* | *p__Firmicutes g__[Ruminococcus] gauvreauii group* | *p__Bacteroidetes g__Odoribacter* |
| *p__Bacteroidetes g__Prevotella 9* | *p__Bacteroidetes g__Butyricimonas* | *p__Proteobacteria f__Burkholderiaceae* |
| *p__Firmicutes g__Coprococcus 3* | *p__Firmicutes g__Clostridium sensu stricto 1* | *p__Proteobacteria g__Massilia* |
| *p__Firmicutes g__Clostridium sensu stricto 1* | *p__Firmicutes g__Lachnospiraceae NK4A136 group* | *p__Proteobacteria f__Pasteurellaceae* |
| *p__Firmicutes g__Ruminococcus 1* | *p__Firmicutes g__Butyricicoccus* | *p__Proteobacteria g__Neisseria* |
| *p__Firmicutes g__Ruminococcaceae UCG-013* | *p__Firmicutes g__Lachnospiraceae UCG-004* | *p__Proteobacteria g__Actinobacillus* |
| *p__Firmicutes g__Roseburia* | *p__Firmicutes f__Ruminococcaceae* | *p__Bacteroidetes g__Rikenellaceae RC9 gut group* |
| *p__Firmicutes g__Lactobacillus* | *p__Firmicutes g__Lachnospiraceae ND3007 group* | *p__Fusobacteria g__Leptotrichia* |
| *p__Firmicutes g__Dialister* | *p__Firmicutes g__Lachnospiraceae FCS020 group* | *p__Bacteroidetes g__Prevotellaceae NK3B31 group* |
| *p__Firmicutes f__Ruminococcaceae* | *p__Firmicutes g__Peptoniphilus* | *p__Proteobacteria g__Methylobacterium* |
| *p__Firmicutes g__Phascolarctobacterium* | *p__Firmicutes g__Oribacterium* | *p__Proteobacteria g__Paracoccus* |
| *p__Firmicutes g__Ruminococcaceae UCG-002* | *p__Firmicutes g__Anaerococcus* | *p__Proteobacteria g__Paucibacter* |
| *p__Firmicutes g__[Ruminococcus] gnavus group* | *p__Firmicutes g__Lachnospiraceae UCG-010* | *p__Proteobacteria g__gut metagenome* |
| *p__Firmicutes g__Ruminococcaceae UCG-004* | *p__Firmicutes g__Acidaminococcus* | *p__Firmicutes g__Oscillibacter* |
| *p__Firmicutes g__Coprococcus 2* | *p__Firmicutes g__[Eubacterium] eligens group* | *p__Proteobacteria g__Desulfovibrio* |
| *p__Firmicutes g__Ruminococcaceae UCG-014* | *p__Firmicutes g__[Eubacterium] nodatum group* | *p__Proteobacteria g__Pseudomonas* |
| *p__Firmicutes g__Butyricicoccus* | *p__Firmicutes g__Megasphaera* | *p__Proteobacteria g__Parasutterella* |
| *p__Bacteroidetes g__Prevotella 2* | *p__Firmicutes g__Lachnospira* | *p__Proteobacteria g__Acinetobacter* |
| *p__Firmicutes g__Coprococcus 1* | *p__Firmicutes g__Moryella* | *p__Proteobacteria g__Sphingobium* |
| *p__Firmicutes g__Lachnospiraceae NK4A136 group* | *p__Firmicutes g__Lachnospiraceae UCG-008* | *p__Firmicutes g__Peptococcus* |
| *p__Actinobacteria g__Bifidobacterium* | *p__Firmicutes g__Coprococcus 2* | *p__Proteobacteria g__Klebsiella* |
| *p__Firmicutes g__Lachnoclostridium* | *p__Firmicutes g__Oscillospira* | *p__Proteobacteria c__Alphaproteobacteria* |
| *p__Firmicutes g__Marvinbryantia* | *p__Firmicutes g__Butyrivibrio* | *p__Bacteroidetes g__Porphyromonas* |
| *p__Firmicutes g__Ruminococcaceae UCG-005* | *p__Firmicutes g__Eubacterium* | *p__Firmicutes g__Enterococcus* |
| *p__Firmicutes g__Moryella* | *p__Firmicutes g__[Eubacterium] saphenum group* | *p__Proteobacteria g__Aggregatibacter* |
| *p__Firmicutes g__Lachnospiraceae FCS020 group* | *p__Firmicutes g__Lachnospiraceae UCG-001* | *p__Proteobacteria g__Burkholderia-Caballeronia-Paraburkholderia* |
| *p__Firmicutes g__Ruminococcaceae NK4A214 group* | *p__Firmicutes g__Anaerotruncus* | *p__Proteobacteria g__Bilophila* |
| *p__Firmicutes g__Lachnospiraceae ND3007 group* | *p__Firmicutes g__Shuttleworthia* | *p__Cyanobacteria o__Gastranaerophilales* |
| *p__Firmicutes g__Negativibacillus* | *p__Firmicutes g__Catonella* | *p__Proteobacteria f__uncultured* |
| *p__Firmicutes g__Veillonella* | *p__Firmicutes g__Lachnoclostridium 5* | *p__Proteobacteria g__Mesorhizobium* |
| *p__Bacteroidetes g__Butyricimonas* | *p__Firmicutes g__Anaerofustis* | *p__Epsilonbacteraeota g__Helicobacter* |
| *p__Firmicutes g__Eubacterium* | *p__Firmicutes g__Lachnospiraceae UCG-002* | *p__Epsilonbacteraeota g__Campylobacter* |

| | | |
|---|---|---|
| p__Bacteroidetes g__Prevotella 7 | p__Firmicutes g__Lachnospiraceae AC2044 group | p__Proteobacteria g__Brevundimonas |
| p__Firmicutes g__Ruminococcaceae UCG-003 | p__Firmicutes g__Lachnospiraceae NK4B4 group | p__Proteobacteria g__Devosia |
| p__Firmicutes g__[Eubacterium] eligens group | p__Firmicutes g__Lachnospiraceae FE2018 group | p__Proteobacteria g__Reyranella |
| p__Firmicutes g__Lachnospiraceae UCG-001 | p__Firmicutes g__Lachnospiraceae NC2004 group | p__Proteobacteria g__Novosphingobium |
| p__Firmicutes g__Ruminococcaceae UCG-010 | p__Firmicutes g__Lachnospiraceae NK3A20 group | p__Proteobacteria g__Morganella |
| p__Firmicutes g__Lachnospiraceae NK4B4 group | | p__Proteobacteria g__uncultured bacterium |
| p__Firmicutes g__Acidaminococcus | | p__Proteobacteria o__Aeromonadales |
| p__Firmicutes o__Lactobacillales | | p__Proteobacteria f__Beijerinckiaceae |
| p__Firmicutes g__Megasphaera | | p__Proteobacteria g__DSSD61 |
| p__Firmicutes g__[Eubacterium] nodatum group | | p__Proteobacteria o__Betaproteobacteriales |
| p__Firmicutes g__Megamonas | | p__Proteobacteria g__uncultured |
| p__Firmicutes g__Lachnospiraceae UCG-008 | | p__Spirochaetes g__Treponema 2 |
| p__Firmicutes g__Lachnospira | | p__Fusobacteria g__uncultured |
| p__Firmicutes g__Oscillospira | | p__Proteobacteria g__Eikenella |
| p__Verrucomicrobia g__Akkermansia | | p__Proteobacteria g__Phreatobacter |
| p__Firmicutes g__Butyrivibrio | | p__Proteobacteria f__Sphingomonadaceae |
| p__Firmicutes g__Lachnospiraceae UCG-004 | | p__Proteobacteria f__Rhodobacteraceae |
| p__Firmicutes g__Lachnospiraceae NK3A20 group | | p__Proteobacteria g__Sphingopyxis |
| p__Firmicutes g__Ruminococcaceae UCG-009 | | p__Proteobacteria g__Sphingomonas |
| p__Firmicutes g__Anaerofustis | | p__Bacteroidetes g__Prevotellaceae UCG-001 |
| p__Firmicutes g__Lachnospiraceae UCG-002 | | p__Proteobacteria g__Oxalobacter |
| p__Firmicutes g__Anaerotruncus | | p__Proteobacteria g__Blastomonas |
| p__Firmicutes g__Ruminococcaceae UCG-007 | | p__Actinobacteria g__Mycobacterium |
| p__Firmicutes g__Lachnospiraceae FE2018 group | | p__Proteobacteria f__Xanthobacteraceae |
| p__Firmicutes g__Lachnospiraceae UCG-010 | | p__Proteobacteria g__Acidovorax |
| p__Firmicutes g__Peptoniphilus | | p__Proteobacteria g__Bradyrhizobium |
| p__Bacteroidetes g__Prevotella | | p__Fusobacteria o__Fusobacteriales |
| p__Firmicutes g__Ruminococcaceae UCG-008 | | p__Proteobacteria g__Acidibacter |
| p__Firmicutes g__Anaerococcus | | p__Fusobacteria g__Sneathia |
| p__Firmicutes g__Oribacterium | | p__Proteobacteria g__Aeromonas |
| p__Proteobacteria g__Succinivibrio | | p__Proteobacteria g__Cardiobacterium |
| p__Firmicutes g__Lachnospiraceae NC2004 group | | p__Proteobacteria g__Schlegelella |
| p__Actinobacteria f__Propionibacteriaceae | | p__Proteobacteria g__Roseomonas |
| p__Bacteroidetes g__Prevotella 6 | | p__Firmicutes g__Bacillus |
| p__Firmicutes g__[Eubacterium] saphenum group | | p__Proteobacteria g__Bosea |
| p__Firmicutes g__Catonella | | p__Proteobacteria g__Legionella |
| p__Firmicutes g__Lachnoclostridium 5 | | p__Proteobacteria g__1174-901-12 |
| p__Firmicutes g__Lachnospiraceae AC2044 group | | p__Proteobacteria g__Phyllobacterium |
| p__Firmicutes g__Selenomonas | | p__Firmicutes g__Mogibacterium |
| p__Firmicutes g__Shuttleworthia | | p__Proteobacteria g__Thiothrix |
| | | p__Proteobacteria g__Methylophilus |
| | | p__Proteobacteria |
| | | p__Proteobacteria g__Ralstonia |
| | | p__Proteobacteria g__Belnapia |
| | | p__Proteobacteria g__Caulobacter |
| | | p__Proteobacteria g__Skermanella |

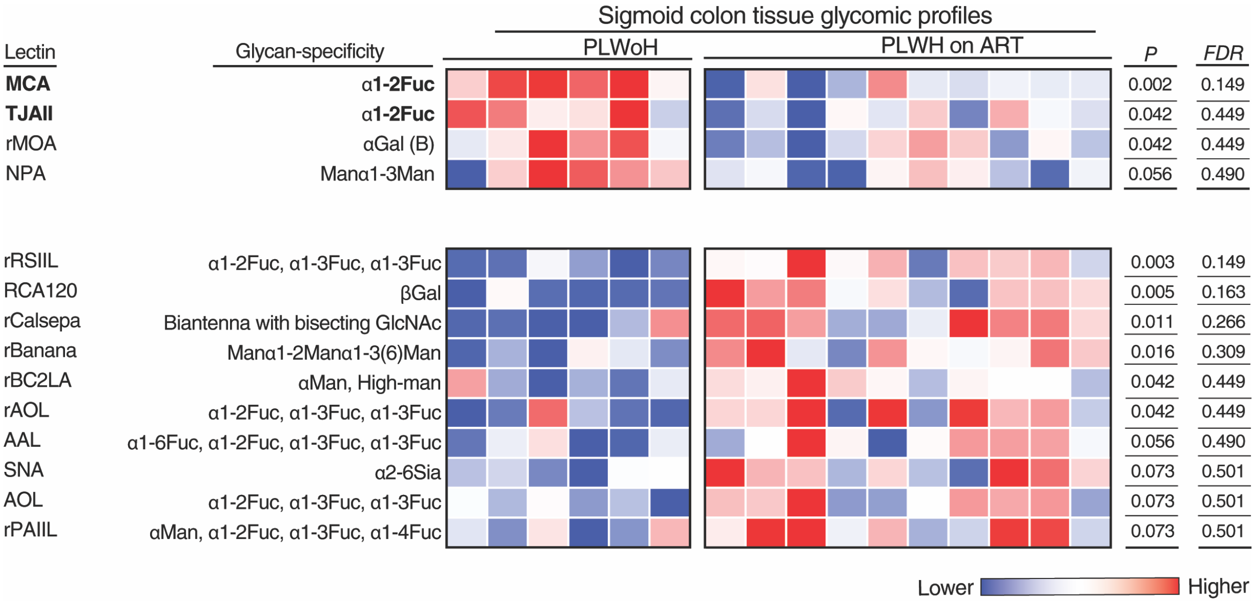


**Figure S1. Sigmoid colon biopsies from PLWH on ART exhibit lower α1,2-fucose than biopsies from PLWoH, related to Figure 1.** Heatmap depicting the relative levels of multiple glycans measured by lectin microarray in sigmoid colon tissues from PLWH on ART and PLWoH controls. Red indicates higher expression, and blue indicates lower expression. *P* values were calculated using the Mann-Whitney U test, and false discovery rates were calculated using the Benjamini-Hochberg method. Each column represents one sample from one individual.

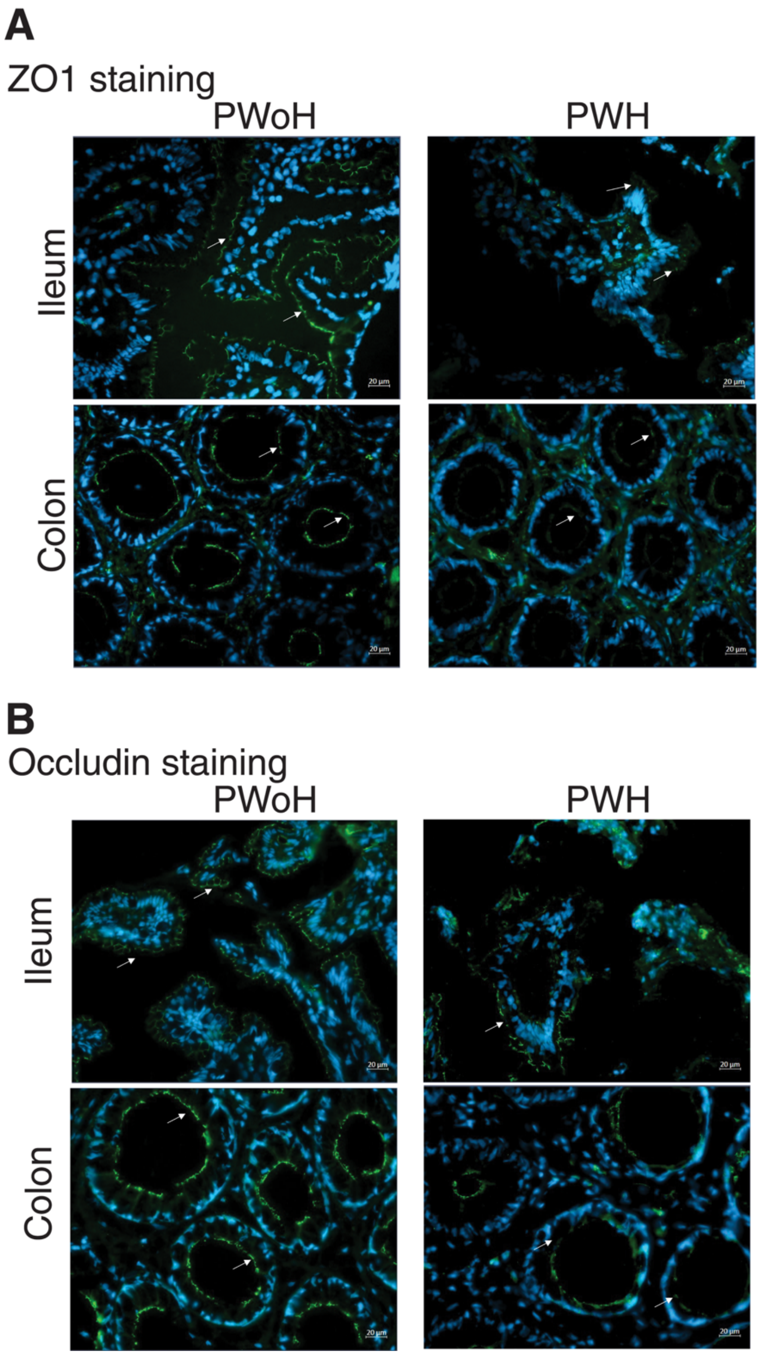


**Figure S2. Immunofluorescence assessment of tight junction proteins in ileal and colonic tissues, related to Figure 5.** Ileum and colon samples from PWoH and PLWH on ART were stained for ZO-1 (A) or occludin (B) expression (green). Nuclei were stained with DAPI (blue). Images, captured at 40x magnification on a Zeiss Axio Observer 7 microscope, have a scale bar of 20 µm.

### Inflammatory signaling

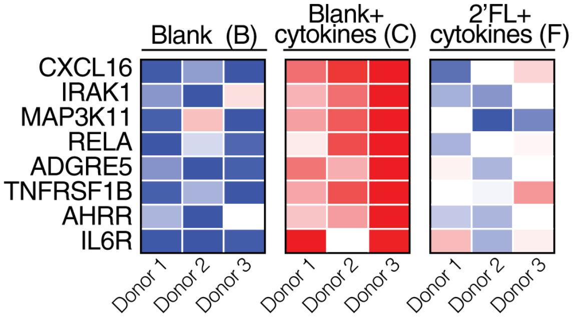


| Gene | *P* (C vs. B) | *FDR* (C vs. B) | *P* (F vs. B) | *FDR* (F vs. B) | *P* (F vs. C) |
|---|---|---|---|---|---|
| CXCL16 | <0.0001 | 0.02 | 0.17 | 1 | 0.007 |
| IRAK1 | 0.007 | 0.21 | 0.75 | 1 | 0.018 |
| MAP3K11 | 0.001 | 0.087 | 0.67 | 1 | <0.0001 |
| RELA | <0.0001 | 0.015 | 0.07 | 1 | 0.019 |
| ADGRE5 | <0.0001 | 0.014 | 0.09 | 1 | 0.011 |
| TNFRSF1B | <0.0001 | 0.007 | 0.03 | 1 | 0.026 |
| AHRR | <0.0001 | 0.015 | 0.44 | 1 | 0.001 |
| IL6R | <0.0001 | 0.001 | 0.001 | 0.4 | 0.076 |

Lower — Higher
Expression

### Epithelial stress and homeostasis

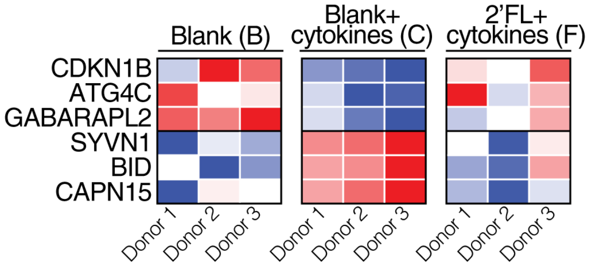


| Gene | *P* (C vs. B) | *FDR* (C vs. B) | *P* (F vs. B) | *FDR* (F vs. B) | *P* (F vs. C) |
|---|---|---|---|---|---|
| CDKN1B | <0.0001 | 0.02 | 0.64 | 1 | 0.0003 |
| ATG4C | 0.004 | 0.17 | 0.84 | 1 | 0.002 |
| GABARAPL2 | 0.0001 | 0.04 | 0.32 | 1 | 0.005 |
| SYVN1 | <0.0001 | 0.006 | 0.67 | 1 | <0.0001 |
| BID | <0.0001 | 0.03 | 0.36 | 1 | 0.003 |
| CAPN15 | 0.0002 | 0.047 | 0.82 | 1 | <0.0001 |

Lower — Higher
Expression

### Epithelial architecture and polarity

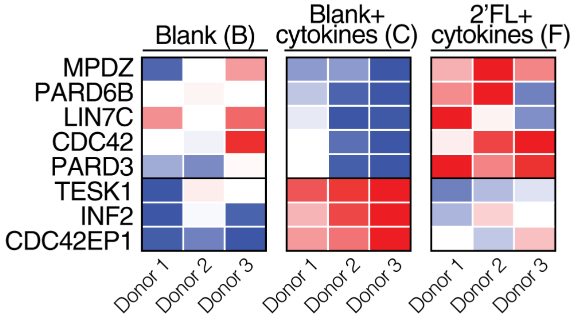


| Gene | *P* (C vs. B) | *FDR* (C vs. B) | *P* (F vs. B) | *FDR* (F vs. B) | *P* (F vs. C) |
|---|---|---|---|---|---|
| MPDZ | 0.22 | NA | 0.05 | 1 | 0.001 |
| PARD6B | 0.005 | 0.17 | 0.75 | 1 | 0.012 |
| LIN7C | 0.017 | 0.27 | 0.67 | 1 | 0.048 |
| CDC42 | 0.06 | 0.40 | 0.77 | 1 | 0.030 |
| PARD3 | 0.51 | 0.79 | 0.01 | 1 | 0.001 |
| TESK1 | 0.0002 | 0.05 | 0.93 | 1 | 0.0001 |
| INF2 | 0.0001 | 0.04 | 0.26 | 1 | 0.007 |
| CDC42EP1 | <0.0001 | 0.001 | 0.01 | 1 | 0.014 |

Lower — Higher
Expression

### Mucosal repair, remodeling, and metabolism

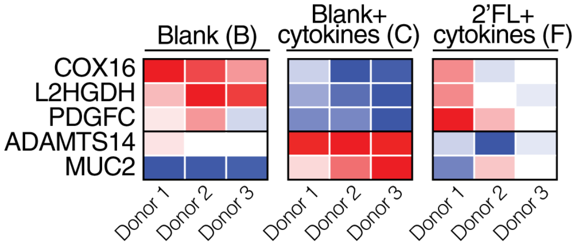


| Gene | *P* (C vs. B) | *FDR* (C vs. B) | *P* (F vs. B) | *FDR* (F vs. B) | *P* (F vs. C) |
|---|---|---|---|---|---|
| COX16 | 0.003 | 0.05 | 0.26 | 1 | 0.01 |
| L2HGDH | <0.0001 | 0.004 | 0.17 | 1 | 0.001 |
| PDGFC | 0.001 | 0.099 | 0.47 | 1 | <0.0001 |
| ADAMTS14 | 0.010 | 0.23 | 0.17 | 1 | 0.0001 |
| MUC2 | <0.0001 | 0.002 | 0.01 | 1 | 0.02 |

Lower — Higher

**Figure S3. 2′FL-containing fermentation supernatants shift cytokine-exposed colon organoids toward a gut-protective transcriptional state, related to Figure 7.** Heatmaps show normalized expression of selected gut-relevant genes in colon organoids treated with blank fermentation supernatant Blank, B, blank fermentation supernatant plus cytokines Blank + cytokines, C, or 2′FL-containing fermentation supernatant plus cytokines 2′FL + cytokines, F across three independent donors. P values were calculated using a paired Wald test in DESeq2. FDR-adjusted P values were calculated using the Benjamini-Hochberg procedure.
